# A new method for determining ribosomal DNA copy number shows differences between *Saccharomyces cerevisiae* populations

**DOI:** 10.1101/2021.01.21.427686

**Authors:** Diksha Sharma, Sylvie Hermann-Le Denmat, Nicholas J. Matzke, Katherine Hannan, Ross D. Hannan, Justin M. O’Sullivan, Austen R. D. Ganley

## Abstract

Ribosomal DNA genes (rDNA) encode the major ribosomal RNAs (rRNA) and in eukaryotic genomes are typically present as one or more arrays of tandem repeats. Species have characteristic rDNA copy numbers, ranging from tens to thousands of copies, with the number thought to be redundant for rRNA production. However, the tandem rDNA repeats are prone to recombination-mediated changes in copy number, resulting in substantial intra-species copy number variation. There is growing evidence that these copy number differences can have phenotypic consequences. However, we lack a comprehensive understanding of what determines rDNA copy number, how it evolves, and what the consequences are, in part because of difficulties in quantifying copy number. Here, we developed a genomic sequence read approach that estimates rDNA copy number from the modal coverage of the rDNA and whole genome to help overcome limitations in quantifying copy number with existing mean coverage-based approaches. We validated our method using strains of the yeast *Saccharomyces cerevisiae* with previously-determined rDNA copy numbers, and then applied our pipeline to investigate rDNA copy number in a global sample of 788 yeast isolates. We found that wild yeast have a mean copy number of 92, consistent with what is reported for other fungi but much lower than in laboratory strains. We also show that different populations have different rDNA copy numbers. These differences can partially be explained by phylogeny, but other factors such as environment are also likely to contribute to population differences in copy number. Our results demonstrate the utility of the modal coverage method, and highlight the high level of rDNA copy number variation within and between populations.

**Author summary:** The ribosomal RNA gene repeats (rDNA) form large tandem repeat arrays in most eukaryote genomes. Their tandem arrangement makes the rDNA prone to copy number variation, and there is increasing evidence that this copy number variation has phenotypic consequences. However, difficulties in measuring rDNA copy number hamper investigation into rDNA copy number dynamics and their significance. Here we developed a novel bioinformatics method for measuring rDNA copy number from whole genome sequence data that is based on the modal sequence read coverage. We established parameters for optimal performance of the method and validated it using yeast strains of known rDNA copy numbers. We then applied the method to a dataset of almost 800 global yeast isolates and demonstrate that yeast populations have different rDNA copy numbers that partially correlate with phylogeny. Our work provides a simple and accurate method for determining rDNA copy number that leverages the growing number of whole genome datasets, and highlights the dynamic nature of rDNA copy number.

## Introduction

The ribosomal RNA gene repeats (rDNA) encode the major ribosomal RNA (rRNA) components of the ribosome, and thus are essential for ribosome biogenesis and protein translation. In most eukaryotes the rDNA forms large tandem repeat arrays on one or more chromosomes [1]. Each repeat unit comprises a coding region transcribed by RNA polymerase I (Pol-I) that encodes 18S, 5.8S and 28S rRNA [2], and an intergenic spacer region (IGS) that separates adjacent coding regions (**Fig 1**). The number of rDNA repeat copies varies widely between species, typically from tens to hundreds of thousands of copies [1, 3–5]. However, each species appears to have a ‘set’ or homeostatic (in the sense of [6]) rDNA copy number that is returned to if the number of copies deviates [7–10]. Deviation in rDNA copy number between individuals within a species is well documented and can be substantial [11–16]. This copy number variation is thought to be tolerated because of redundancy in rDNA copies [8, 17]. This redundancy can partly be explained by the striking observation that only a subset of the repeats is transcribed at any one time [2]. Thus, cells can compensate for changes in rDNA copy number by activating or silencing repeats to maintain the same transcriptional output [18]. The variation in rDNA copy number is a consequence of unequal homologous recombination, which results in loss or gain of rDNA copies [8, 19–22]. This copy number variation is, somewhat counter-intuitively, what drives the high levels of sequence homogeneity observed between the rDNA copies within a genome, a pattern known as concerted evolution [23–25]. Recent results in *Saccharomyces cerevisiae* revealed an elegant mechanism through which homeostatic rDNA copy number is maintained in the face of rDNA copy number change via the abundance of the Pol-I transcription factor UAF (upstream activating factor) and the histone deacetylase Sir2 [26]. However, the selective pressure(s) that determines what the homeostatic rDNA copy number is remains unknown. Nevertheless, there is growing evidence that rDNA copy number and the proportion of active/silent rDNA copies impact several aspects of cell biology beyond simply rRNA production [8, 12, 17, 22, 27–35].

**Figure 1.**
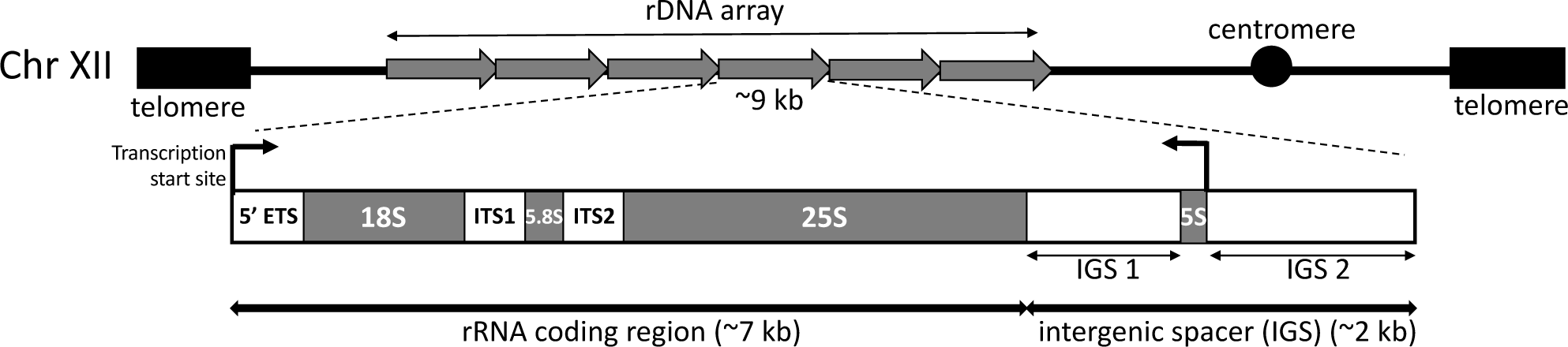
Organization of the rDNA repeats in *Saccharomyces cerevisiae*. Top shows a schematic of tandemly-repeated units in the rDNA array located on chromosome XII. Bottom shows the organization of an individual rDNA repeat including transcription start sites, the 5’ external transcribed spacer (5’ETS), the rRNA (18S, 5.8S and 28S) coding genes, the two internal transcribed spacers (ITS1 and 2), and the intergenic spacer (IGS). The IGS is divided into two by a 5S rRNA gene. Schematic is not to scale.

Interest in the phenotypic consequences of rDNA copy number variation has led to a number of approaches being used to measure it. These include molecular biology approaches such as quantitative DNA hybridization [36–39], pulsed field gel electrophoresis (PFGE) [40, 41], quantitative real-time PCR (qPCR) [15, 42–46] and digital droplet PCR (ddPCR) [47, 48]. A major advance in the measurement of rDNA copy number has been the emergence of bioinformatic approaches that use whole genome (WG) next generation sequencing (NGS) reads to estimate copy number, based on the rationale that sequence coverage of the rDNA correlates with copy number. This correlation is a consequence of concerted evolution, with the high sequence identity between repeats resulting in reads from all rDNA copies mapping to a single reference rDNA unit, thus providing a high coverage signal that is proportional to copy number. Existing bioinformatic approaches calculate the mean rDNA read coverage and normalize to the mean WG coverage to estimate copy number [5, 12, 25, 34, 49], thus assuming that mean coverage represents the “true coverage” for both the rDNA and the WG. However, there are reasons to suspect this mean coverage approach assumption might not always hold. Repetitive elements (e.g. microsatellites and transposons), PCR/sequencing bias (which is particularly evident for the rDNA [50–54]; **Supplementary Figure 1**), and large-scale mutations such as aneuploidies and segmental duplications may all cause the measured mean coverage to differ from the real coverage. While efforts have been made to address some of these potential confounders [12, 55, 56], estimated copy number varies depending on which region of the rDNA is used [12, 34], thus the accuracy of this mean read coverage approach has been called into question [5, 46].

Here we present a bioinformatics pipeline that measures rDNA copy number using modal (most frequent) NGS read coverage as a way to overcome the limitations of the mean coverage bioinformatics approach. We assessed the parameters important for performance and validated the pipeline using *S. cerevisiae* strains with known rDNA copy numbers. We then employed our pipeline to investigate whether *S. cerevisiae* populations maintain different homeostatic rDNA copy numbers.

## Materials and Methods

### Modified Saccharomyces cerevisiae genome

Chromosome sequences for *S. cerevisiae* strain W303 were obtained from the NCBI (accession CM001806.1 - CM001823.1) and concatenated. rDNA copies present within the W303 reference genome were identified using BLAST and removed using Geneious (v. 11.0.3). The *S. cerevisiae* W303 strain rDNA repeat unit from [23] was then added as an extrachromosomal rDNA reference, and this modified W303 yeast reference genome (W303-rDNA) was used in subsequent analyses.

### Yeast strains/isolates and growth conditions

Yeast strains/isolates that were cultured are listed in **Table 1**. Culturing was performed in liquid or solid (2% agar) YPD (1% w/v yeast extract, 2% w/v peptone and 2 % w/v D+ glucose) medium at 30°C.

**Table 1:**
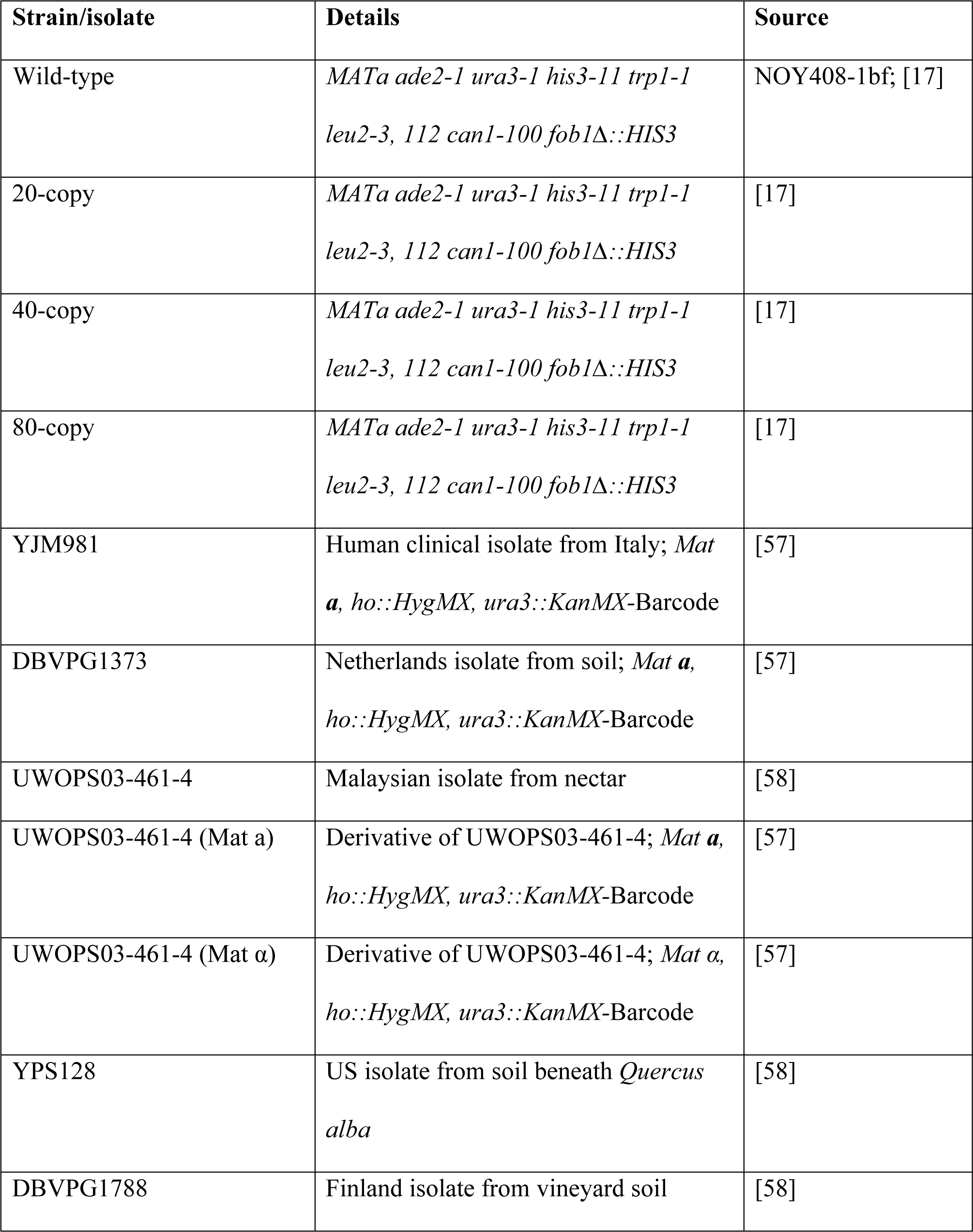
*S. cerevisiae* strains/isolates cultured in this study.

### Genomic DNA extraction

High molecular weight genomic DNA (gDNA) was isolated as follows. Cell pellets from 3–5 mL liquid cultures were washed in 500 µL of 50 mM EDTA pH 8 and resuspended in 200 µL of 50 mM EDTA pH 8 supplemented with zymolyase (3 mg/mL). After 1 hr at 37°C, the cell lysate was mixed with 20 µL of 10% sodium dodecyl sulfate then with 150 µL of 3 M potassium acetate (KAc) and incubated on ice for 1 hr. 100 µL of phenol-chloroform-isoamyl alcohol was added to the SDS-KAc suspension, and, following vortexing and centrifugation, 600 µL of propanol-2 were added to the aqueous supernatant (≈ 300 µL). The nucleic acid pellet was washed three times in 70% EtOH, dried and resuspended in PCR grade water supplemented with RNase A (0.3 mg/mL). After 1 hr at 37°C, samples were stored at −20°C.

### Whole genome sequence data

gDNA extracted from four isogenic strains with different rDNA copy numbers (WT, 20-copy, 40-copy and 80-copy; **Table 1**) was sequenced using Illumina MiSeq (**Supplementary Table 1**). The raw sequence files are available through the NCBI SRA (accession number SUB7882611).

### Read preparation

Paired-end reads were combined and quality checked using SolexaQA [59]. Low-quality ends of reads (score cutoff 13) were trimmed using DynamicTrim, and short reads were removed using a length cutoff of 50 bp with LengthSort, both within SolexaQA, as follows:

**command:** ∼/path/to/solexaQA/SolexaQA++ dynamictrim /fastq/file

**command:** ∼/path/to/solexaQA/SolexaQA++ lengthsort -l 50 /trimmed/fastq/file

### Obtaining whole genome and rDNA coverages

The W303-rDNA reference genome was indexed using bowtie2 (v. 2.3.2):

**command:** ∼/bowtie2-2.3.2/bowtie2-build <reference_in> <bt2_base>

Coverage files for the whole genome and rDNA were obtained using a four step pipeline:

**Step-1:** Processed reads were mapped to the indexed W303-rDNA genome using bowtie2:

**command:** ∼/bowtie2-2.3.2/bowtie2 -x /path/to/indexed/genome/ -U /path/to/trimmed/reads/ -S /output SAM file/

**Step-2:** The subsequent SAM format alignment was converted to BAM format using SAMtools (v. 1.8):

**command:** ∼/samtools-1.8/samtools view -b -S -o <output_BAM> <input_SAM>

**Step-3:** Mapped reads in the BAM file were sorted according to the location they mapped to in W303-rDNA using SAMtools:

**command:** ∼/samtools-1.8/samtools sort <input_BAM> -o <output_sorted.bam>

**Step-4:** Per-base read coverages across the entire W303-rDNA genome and the rDNA were obtained using BEDtools (v. 2.26.0):

**command:** ∼/bedtools genomecov -ibam <aligned_sorted.bam> -g <reference_genome.fasta> -d <bedtools_coverage_WG.txt>

**command:** grep “rDNA_BLAST” <bedtools_coverage_WG.txt> <rDNA_bedtools_coverage.txt>

### Calculation of rDNA copy number using modal coverage

Coverage frequency tables for the rDNA and whole genome (excluding mitochondrial DNA and plasmids) were obtained from per-base read coverage files by computing the mean coverage over a given sliding window size with a slide of 1 bp. The mean coverage for each sliding window was then allocated into a coverage bin. The bin that includes read coverage of zero was subsequently removed. The three highest frequency coverage bins from both the rDNA and whole genome frequency tables were used to calculate rDNA copy number as follows:

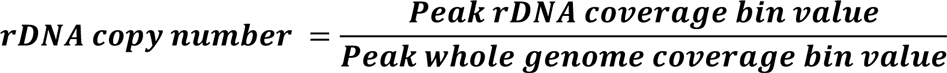

rDNA copy number estimates are the mean of all pairwise combinations of these copy number values (**Supplementary Figure 2**).

### Pipeline availability

The pipeline for modal calculation of rDNA copy number from an alignment of sequence reads to a reference genome containing one rDNA copy is available through Github (https://github.com/diksha1621/rDNA-copy-number-pipeline).

### Calculation of rDNA copy number using mean and median coverage

Per-base read coverage across W303-rDNA from Bedtools was input into custom R-scripts to obtain mean and median coverage values for the whole genome and rDNA after removing the rDNA, 2-micron plasmid, and mitochondrial DNA coverage values from the whole genome calculation. rDNA copy number was then calculated for the mean and median data as follows:

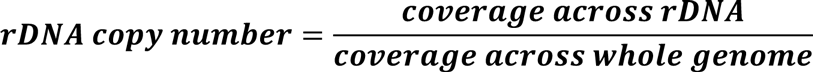

### Subsampling

To generate different coverage levels for copy number estimation, sequence reads were randomly downsampled using the seqtk tool (https://github.com/lh3/seqtk):

**command:** ∼/seqtk/seqtk sample –s$RANDOM <name of fastqfile> <number of reads required> <outputfile>

### rDNA copy number measurement by ddPCR

At least three independent cultures (biological replicates) were generated for each isolate using one independent colony per culture. To evaluate rDNA copy number variation over generations, cultures were propagated over four days (∼60 generations) as follows: individual colonies were initially grown in 3 mL YPD for 24 hr. 30 µL of this was used to inoculate 3 mL YPD and this was grown for another 24 hr. This process was repeated for four days. Cells were harvested after 24 hr (∼15 generations) and four days, and cell pellets frozen at −80°C. gDNA was extracted as above, then linearized by *Xba*I in NEB2 buffer following the manufacturer’s instructions (NEB) to individualize rDNA repeats. gDNA linearization was verified by separation on agarose gels and DNA concentration measured on a Qubit Fluorometer using the Qubit dsDNA HS assay (Thermo Fisher). Linearized gDNA was brought to 2 pg/µL by serial dilution. EvaGreen master mixes were prepared with an rDNA primer pair (rDNAScSp_F2 5’-ATCTCTTGGTTCTCGCATCG-3’, rDNAScSp_R2 5’-GGAAATGACGCTCAAACAGG-3’) or a single copy *RPS3* gene primer pair (RPS3ScSp_F2 5’-CACTCCAACCAAGACCGAAG-3’, RPS3ScSp_R2 5’-GACAAACCACGGTCTTGAAC-3’). *RPS3* and rDNA ddPCR reactions were performed with 2 µL (4 pg) of the same linearized gDNA dilution as template. Droplet generation and endpoint PCR were performed following the manufacturer’s instructions, and droplets were read using a QX200 droplet reader (BioRad). Quantification was performed using QuantaSoft Analysis Pro (v. 1.0.596). rDNA copy number was determined by the (rDNA copy/µL)/(*RPS3* copy/µL) ratio.

### Pulse field gel electrophoresis (PFGE)

To make chromosome plugs [21], cells from overnight liquid cultures were resuspended in 50 mM EDTA pH 8.0 to 2.10^9^ cells/mL, transferred to 45°C, and mixed with an equal volume of 1.5% low melting point agarose in 50 mM EDTA prewarmed to 45°C. The mixture was transferred by gentle pipetting to PFGE plug molds (BioRad) to set at 4°C for 15 min. Plugs were transferred to fresh spheroplasting solution (1 M Sorbitol, 20 mM EDTA pH 8.0, 10 mM Tris-HCl pH 7.5, 14 mM 2-mercaptoethanol, 2 mg/mL zymolyase). After 6 hr incubation at 37°C with occasional inversion, plugs were washed for 15 min in LDS buffer (1% lithium dodecyl sulphate, 100 mM EDTA pH 8.0, 10 mM Tris-HCl pH 8.0), before overnight incubation at 37°C in the same buffer with gentle shaking. Plugs were incubated twice for 30 min each in NDS buffer (500 mM EDTA, 10 mM Tris-HCl, 1% sarkosyl, pH 9.5) and at least three times for 30 min in TE (10 mM Tris-HCl pH 8.0, 1 mM EDTA pH 8.0). Plugs were stored at 4°C in fresh TE. For restriction digestion, half plugs were pre-washed for two hours in TE, three times for 20 min each in TE, and three times for 20 min each in 300 µL restriction buffer supplemented with 100 µg/mL BSA, all at room temp. Restriction digestion was performed overnight at the recommended temperature in a total volume of 500 µL containing 100 U of restriction endonuclease. Digested plugs were washed in 50 mM EDTA pH 8.0 and stored at 4°C in 50 mM EDTA pH 8.0 before loading. PFGE was performed using 1% agarose gel in 0.5X TBE (Thermo-Fisher) in a CHEF Master XA 170-3670 system (BioRad) with the following parameters: auto algorithm separation range 5 kb - 2 Mb (angle 120°C, run 6 V/cm, initial switch time 0.22 s, final switch time 3 min 24 s, run time 916 min) at 14°C. DNA was visualized by staining in ethidium bromide (5 µg/mL) and imaging (Gel Doc XR+; BioRad).

### 1002 Yeast Genome project rDNA copy number estimation

Illumina reads from the 1002 Yeast Genomes project were obtained from the European Nucleotide Archive (www.ebi.ac.uk/) under accession number ERP014555. We omitted clades with few members, mosaic clades, and unclustered isolates, giving a total of 788 isolates. Reads were downsampled to 10-fold-coverage using seqtk() and rDNA copy number for each isolate was calculated using W303-rDNA as the reference. Bin sizes of 1/200^th^ of the mean coverage for rDNA and 1/50^th^ for the whole genome, and a window size of 600 bp for both estimates, were used. Violin plots were plotted using the ggplot() package in R.

### Phylogenetic analyses

To create a neighbour-joining phylogeny based on rDNA copy number values, rDNA copy number for each isolate (after removing 30 isolates for which SNP data were not available) was normalized on a 0-1 scale. Normalized values were used to calculate pairwise Euclidean distances between each pair of isolates to generate a distance matrix that was applied to construct a phylogeny via neighbour-joining using MEGA X [60].

Phylocorrelograms of copy number and SNP phylogeny were generated using phylosignal [61] (v.1.3; https://cran.r-project.org/web/packages/phylosignal/index.html). Phylocorrelograms representing no phylogenetic signal (a “white noise” random distribution) and high phylogenetic signal (a character evolving on the SNP tree according to a Brownian motion model) were also generated. For the white noise distribution, data were simulated from a normal distribution with mean and standard deviation matching those of the observed copy number data (mean=92.5, sd=30.8). For the Brownian motion model, we first estimated the ancestral mean (z0=83.2) and the rate parameter (σ2=72557.2) from the observed copy number data using the fitContinuous function from geiger [62] (https://cran.r-project.org/package=geiger). Then, we simulated from these parameters on the SNP tree using fastBM from phytools 0.7 (https://cran.r-project.org/package=phytools). Phylocorrelograms were generated for the observed and two simulated datasets, estimating correlations at a series of 100 phylogenetic distances using 100 bootstrap replicates.

### Comparing intra-species variation in rDNA copy number

Copy number estimates for twelve isolates from the 1002 Yeast Genomes data were randomly drawn 1000 times using a custom bash-script to obtain rDNA copy number ranges.

### Statistical analyses

Statistical analyses were performed in R. Significance was calculated using the Welch *t*-test (*t*-test), the non-parametric Wilcoxon-Mann-Whitney test (wilcox test) or ANOVA, with *p*-values considered statistically significant at *p* < 0.05.

## Results and Discussion

### Establishment of a modal coverage bioinformatics pipeline for estimating rDNA copy number

The abundance of data generated from NGS platforms has led a number of studies to use mean read depth to estimate rDNA copy number [5, 12, 25, 34, 49, 55, 56]. However, repeat elements, sequence biases and large-scale changes like aneuploidies can potentially result in non-normal read coverage distributions where the mean coverage does not accurately represent the true coverage. To overcome these limitations, we developed a novel sequence read-based rDNA copy number calculation approach based on the most frequent (modal) coverage. The rationale for this approach is that modal coverage will provide an estimate of the relative coverage representation of a given region in a genome that is more robust to biases away from normality than the mean or median. The approach allocates coverage across a reference genome into coverage bins, and the ratio of the most frequently occurring coverage bins for the rDNA and the WG is then used to calculate rDNA copy number (per haploid genome). We implemented this modal coverage approach as a simple pipeline to calculate rDNA copy number from mapped sequence reads (**Fig 2**). To help smooth across positions that stochastically vary in coverage, an issue that is particularly prevalent with very low coverage datasets, we used a sliding window approach to calculate coverage. Our straightforward pipeline uses a sorted BAM file of reads aligned to a reference genome for which the position of the rDNA is known (either embedded in the genome or as a separate contig) to calculate copy number.

**Figure 2.**
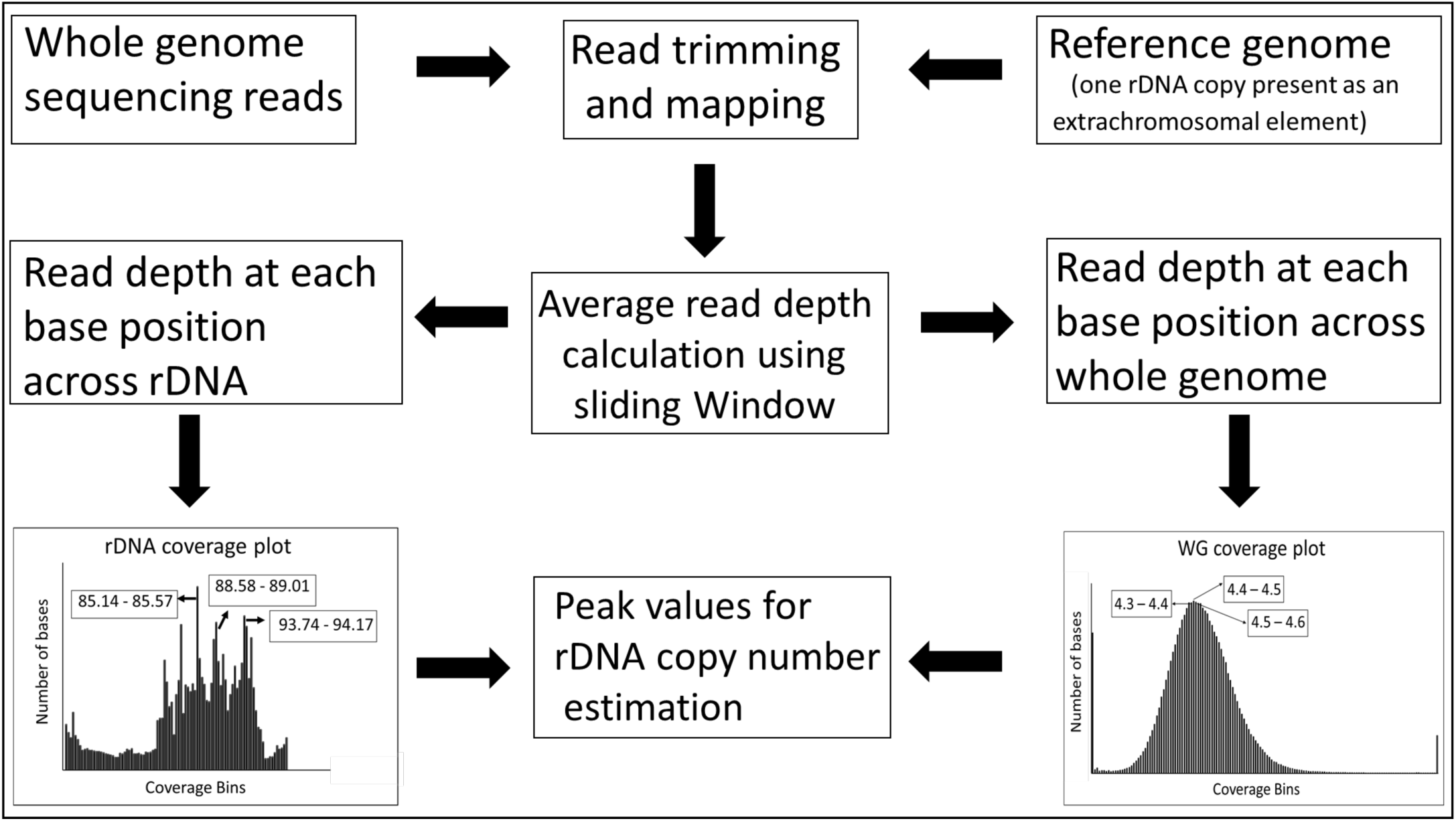
Overview of the modal approach to estimate rDNA copy number from whole genome sequence data. Whole genome (WG) sequence reads are mapped against a reference genome containing a single rDNA copy. Mean read depth for each postion is calculated across the rDNA and the WG using a sliding window, then allocated into coverage bins (shown as histograms). To calculate modal rDNA copy number, the highest frequency coverage bins for both the rDNA and WG are used to compute ratios that represent the rDNA copy number range. The histograms shown were plotted using a 20-copy yeast strain at 5-fold WG coverage with bin sizes of 1/200^th^ of mean coverage for rDNA and 1/50^th^ for WG, and a sliding window of 600 bp for both. The coverage ranges for the three most frequent bins for each are indicated in boxes.

To implement our modal coverage approach, we generated test datasets by performing WG Illumina sequencing of a haploid wild-type laboratory *S. cerevisiae* strain reported to have 150-200 rDNA copies, and three isogenic derivatives in which the rDNA has been artificially reduced to 20, 40 and 80 copies, and “frozen” in place through disruption of a gene (*FOB1*) that promotes rDNA copy number change [17] (**Table 1**). Initially, we investigated which parameters provide the most accurate results by applying our pipeline to the WG sequence data obtained from a strain with 20 rDNA copies (20-copy strain). We obtained a genome-wide read coverage of 13.1-fold (**Supplementary Table 1**) and mapped these reads to the W303-rDNA yeast reference genome that has a single rDNA copy. The mapping output was used to determine per-base coverage values, which were placed into coverage bins using a sliding window. We investigated a range of sliding window sizes, from 100 bp (previously reported to have an approximately normal distribution of WG sequence read coverage [63]) to 1,000 bp (large sliding window sizes, whilst smoothing stochastic coverage variation, converge on the mean coverage as the window size approaches the rDNA unit length). We also assessed the impact of coverage on copy number estimation by downsampling the sequence reads. We ran all these analyses with 100 technical replicates and computed the rDNA copy number means and ranges. We found that, as expected, the accuracy and precision (defined here as <u>similarity to known copy number</u> and <u>copy number range</u>, respectively) of the pipeline was poorer at lower coverage levels, while larger sliding window sizes could compensate for a lack of reads to improve both measures (**Fig 3**). Coverage levels above 10-fold with a sliding window size between 500-800 bp produced accurate rDNA estimates. However, our method also demonstrated adequate performance even with a coverage level of 5-fold, when the sliding window was 600-700 bp (**Fig 3**). We found that the method works similarly when just using the rRNA coding region (**Supplementary Figure 3**) rather than the full repeat, which is important as the full rDNA unit sequence is often not available. We also examined the performance of median coverage, but found that while it had greater precision compared to the modal coverage approach, the accuracy was poorer (**Supplementary Figure 4**). Given the rapid rate at which copy number changes even during vegetative growth [21], the lower precision of our method may more accurately represent the range of copy numbers likely to be present in samples that consist of multiple cells.

**Figure 3.**
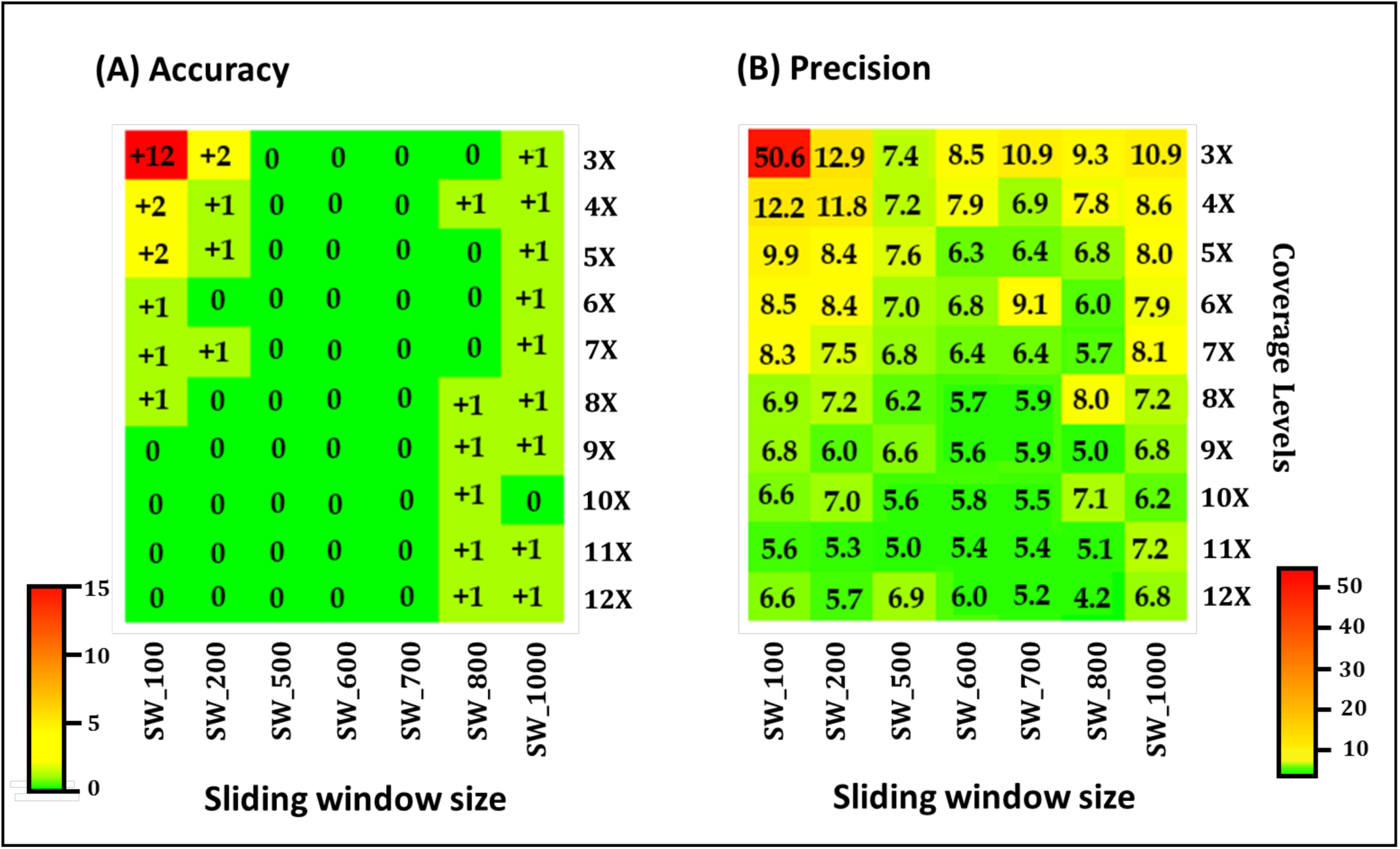
Assessing parameters for rDNA copy number estimation accuracy and precision. Cells represent the (**A**) deviation of the calculated modal rDNA copy number from 20, and (**B**) maximum variation of rDNA copy number calculated from the 100 technical replicates for each coverage level and sliding window (SW) size combination. The heatmap scales used are indicated. In (**A**), rDNA copy number was rounded to the nearest integer.

We then assessed the performance of our pipeline with the 40-copy, 80-copy, and WT *S. cerevisiae* strain data. Illumina WG sequence reads (**Supplementary Table 1**) obtained from these strains were downsampled to generate 100 technical replicates at 10-fold coverage for each strain, and rDNA copy numbers were calculated using our modal coverage pipeline with a sliding window of 600 bp. The resultant rDNA copy numbers were: 32-40 (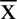= 36 copies) for the 40-copy strain; 57-72 (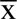= 64 copies) for the 80-copy strain; 129-177 (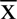= 157 copies) for the WT strain. These values, while similar to the reported copy numbers for these strains, are not identical. Therefore, to check the actual copy numbers of these strains, and to provide a direct validation of our modal pipeline method, we next experimentally determined the rDNA copy numbers of these strains.

We chose ddPCR to experimentally determine rDNA copy number because it is less sensitive than qPCR to biases in secondary structure regions that are common in the rDNA coding region [22]. The ddPCR data showed that the rDNA copy numbers of our strains are similar to those calculated by our modal coverage method, with both methods suggesting that the “80-copy” strain actually has substantially fewer copies than reported (**Supplementary Table 2**; **Supplementary Figure 5A**), perhaps due to a stochastic change in copy number that has occurred in our version of this strain. We also compared our modal coverage approach with the mean coverage calculated from the same datasets. We used a simple mean calculation to match the implementation of our modal approach, using the same down-sampled 10-fold WG coverage datasets. The copy number estimates made using the mean coverage approach were uniformly lower than the other estimates (**Supplementary Table 2**), which we suggest is the result of sequencing biases against regions in the rRNA coding region. Importantly, correlating read coverage and ddPCR copy number estimates showed the modal coverage slope was closer to the expected value of 1 than the mean coverage slope (**Fig 4**). We also estimated the copy number using pulsed field gel electrophoresis based on the size of the restriction fragment encompassing the entire rDNA array divided by the rDNA unit size (accounting for the sizes of the flanking regions), again with consistent results (**Supplementary Figure 5B,C**). Together, these results suggest the modal coverage approach is an accurate way to estimate rDNA copy number.

**Figure 4.**
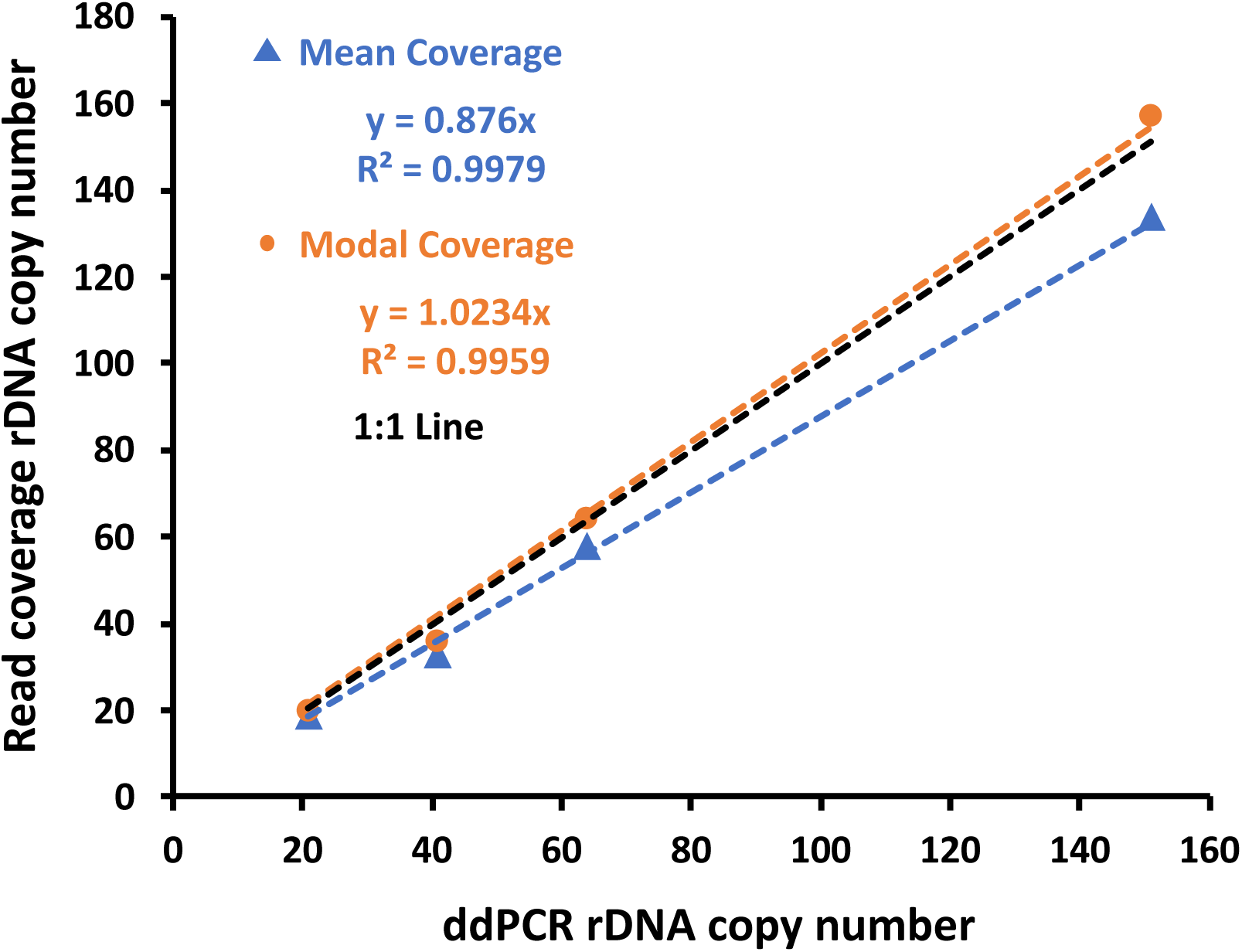
Comparison of modal and mean coverage copy number estimation methods. Plot of rDNA copy number for the 20, 40, 80 and WT *S. cerevisiae* strains (10-fold coverage) calculated using modal (orange line) and mean (blue line) coverage methods versus the copy number determined by ddPCR. The expected 1:1 correlation between read coverage and ddPCR methods is shown in black. Note that while the mean coverage method gives a slightly higher R^2^, the modal coverage results are a closer fit to the expected 1:1 line.

Our results suggest that the modal coverage pipeline provides robust estimates of rDNA copy number even when coverage is less than 5-fold. This reliability may partly be a consequence of the larger sliding window size we used compared to that commonly applied for mean coverage methods. It was previously reported that coverage below ∼65X results in precision issues when estimating rDNA copy number [5]. However, we did not find this, either for our method or when using mean coverage, suggesting the issues might be specific to the approach and/or dataset used in that study. The simple implementation of our modal approach coupled with its good performance make it an attractive method for estimating rDNA copy number from sequence read data. Furthermore, a modal approach is expected to be more robust to features that can perturb mean coverage approaches by skewing coverage distributions, such as repeat elements, large duplications and deletions, regions exhibiting sequencing biases, modest sequence divergence from the reference sequence, and aneuploidies [46]. Although we have developed our pipeline for measuring rDNA copy number, in principle it can be used to calculate copy number for any repeated sequence where all reads map to a single repeat copy and the sequence is known, such as mitochondrial and chloroplast genome copy numbers. Given its strong performance, we applied our method to characterize the inter-population distributions of rDNA copy number in *S. cerevisiae*.

### Within-species evolutionary dynamics of rDNA copy number

Studies in model organisms have provided evidence that each species has a homeostatic copy number which is returned to following copy number perturbations [7–10]. This homeostatic copy number appears to have a genetic basis [5, 26], which suggests it might vary between populations, as well as between species. However, few studies have addressed this question. Given that variation in rDNA copy number has been associated with altered phenotypes [8, 12, 17, 22, 27–35], we decided to undertake a comprehensive assessment of *S. cerevisiae* rDNA copy number at the population level using the global wild yeast dataset from the 1002 Yeast Genomes project [64].

We obtained WG sequence data for 788 isolates from the 1002 Yeast Genomes project. Reads for each isolate were downsampled to 10X genome coverage, mapped to our W303-rDNA reference genome, and rDNA copy numbers estimated using our modal coverage pipeline. The rDNA copy numbers ranged between 22-227 (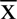= 92) across the 788 isolates (**Supplementary Table 3**). The copy numbers of 11 wild *S. cerevisiae* isolates included in our dataset had previously been estimated [14, 25], and our results are largely consistent with these (**Supplementary Table 4**). However, the copy numbers we estimate are, in general, much lower than those (∼150-200) measured for most laboratory strains (e.g. [17, 38, 41]). We looked to see whether ploidy affects rDNA copy number, given that laboratory strains are predominantly haploid while the wild *S. cerevisiae* isolates we analyzed are mostly diploid. We observed a small difference in copy number between haploid and diploid isolates (104 vs 91 copies, respectively; **Supplementary Figure 6** and **Supplementary Information**), but overall do not find a strong effect of ploidy on copy number. Thus, the copy number differences between lab and most wild *S. cerevisiae* isolates seem to be a property of these isolates.

The difference in copy number between lab and wild *S. cerevisiae* isolates suggests that *S. cerevisiae* populations may harbor different rDNA copy numbers. To test this, we used the 23 phylogenetic clades defined by [64] as proxies for *S. cerevisiae* populations and looked at the distributions of rDNA copy number within and between these populations (**Fig 5**). ANOVA analysis rejects homogeneity of rDNA copy number between these populations (*p* = 4.37e^-15^), suggesting there are population-level differences in *S. cerevisiae* copy number.

**Fig 5.**
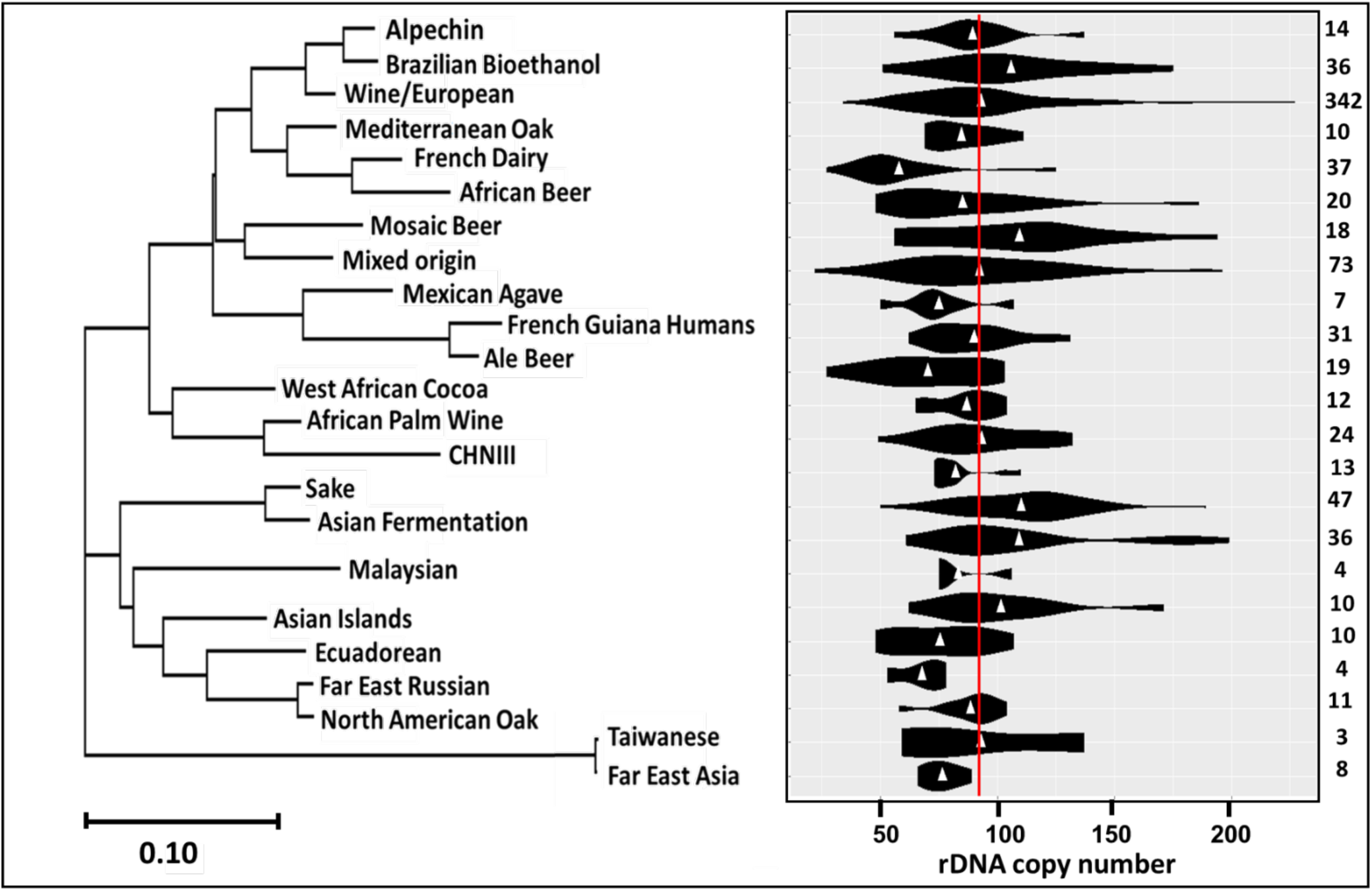
rDNA copy number in *S. cerevisiae* populations. To the left is the phylogeny of the 23 *S. cerevisiae* clades from [64] that encompass the 788 isolates included in this study. The scale represents substitutions per site. On the right, rDNA copy number calculated using the modal coverage method is displayed as a violin plot for each clade with mean population copy numbers indicated by white triangles. Numbers to the right represent the number of isolates in each clade. The red vertical line represents the overall mean rDNA copy number (92 copies). Copy number estimations were determined using 10-fold coverage and a 600 bp sliding window.

We next wanted to look for complementary evidence that *S. cerevisiae* populations have different rDNA copy numbers, as an alternative explanation for our results is different populations happened to have different copy numbers simply due to stochastic variation [21]. If the stochastic variation explanation is correct, we would expect divergent copy numbers to return to a single homeostatic value over time. To test this, we used ddPCR to measure the rDNA copy numbers of six of the 1002 Yeast Genomes project isolates that represent the range of copy numbers observed, including one with three different sub-isolates. We grew three biological replicates of each isolate for ∼60 generations to allow any fluctuation in rDNA copy number to return to the homeostatic level [7]. Despite copy number variation between replicates, which is expected given the high level of stochastic copy number variation, the rDNA copy numbers both before and after the ∼60 generations resemble the copy numbers we estimated from the sequence data and show no tendency to converge on the overall *S. cerevisiae* mean copy number (**Table 2; Supplementary Table 5**). These results strongly suggest that our method of estimating rDNA copy number is robust and that the copy numbers of isolates are not recovering towards a common copy number value. From this we conclude that different *S. cerevisiae* populations have different homeostatic rDNA copy numbers.

**Table 2.** *S. cerevisiae* rDNA copy number does not recover to a common value following ∼60 generations of growth.

| Isolates | rDNA CN at start<sup>a</sup> | rDNA CN after ~60 generations<sup>a</sup> |  | Original modal CN estimation<sup>b</sup> |
| --- | --- | --- | --- | --- |
| *S. cerevisiae* wild-type rep1<sup>c</sup> | 213 | 130 | 185<sup>d</sup> | 157 |
| rep2 |  | 217 |  |  |
| rep3 |  | 208 |  |  |
| YJM981 rep1 | 174 | 120 | 175 | 171 |
| rep2 |  | 183 |  |  |
| rep3 |  | 221 |  |  |
| DBVPG1373 rep1 | 69 | 77 | 85 | 78 |
| rep2 |  | 72 |  |  |
| rep3 |  | 107 |  |  |
| UWOPS03-461-4<sup>c</sup> rep1 | 85 | 113 | 95 | 106 |
| rep2 |  | 88 |  |  |
| rep3 |  | 83 |  |  |
| UWOPS03-461-4<sup>c</sup> (Mata) rep1 | 244 | 164 | 146 |  |
| rep2 |  | 167 |  |  |
| rep3 |  | 106 |  |  |
| UWOPS03-461-4<sup>c</sup> (Matα) rep1 | ND<sup>f</sup> | 108 | 109 |  |
| rep2 |  | 115 |  |  |
| rep3 |  | 105 |  |  |
| YPS128 rep1 | 89 | 87 | 79 | 89 |
| rep2 |  | 73 |  |  |
| rep3 |  | 77 |  |  |
| DBVPG1788 rep1 | 95 | 126 | 108 | 87 |
| rep2 |  | 100 |  |  |
| rep3 |  | 97 |  |  |

<sup>a</sup> Measured using ddPCR <sup>b</sup> Measured using our modal coverage pipeline <sup>c</sup> rep: biological replicate <sup>d</sup> Mean of the three replicates to the nearest integer <sup>e</sup> UWOPS03-461-4 is the parent isolate of UWOPS03-461-4 Mata and UWOPS03-461-4 Mata derivatives <sup>f</sup> Not determined

Copy number has previously been shown to correlate with phylogeny for species across the fungal kingdom [5]. Given the differences in rDNA copy number we observe, we wondered whether a similar correlation exists for *S. cerevisiae* populations. To test this, we constructed a neighbour-joining phylogeny using rDNA copy number as the phylogenetic character for 758 isolates (30 were removed as SNP data were not available) and compared this to the reported *S. cerevisiae* phylogeny created from genomic SNP data [64]. To assess how well the two phylogenies correlate, we used Moran’s Index of spatial autocorrelation *I*, which quantifies the correlation between two traits. Moran’s *I* indicated a small positive correlation between rDNA copy number and phylogeny at short phylogenetic distances (**Fig 6**), but not a significant negative correlation at greater phylogenetic distances like that previously observed above the species level [5]. These results suggest that phylogeny only partially explains the distribution of rDNA copy numbers amongst *S. cerevisiae* populations.

**Figure 6.**
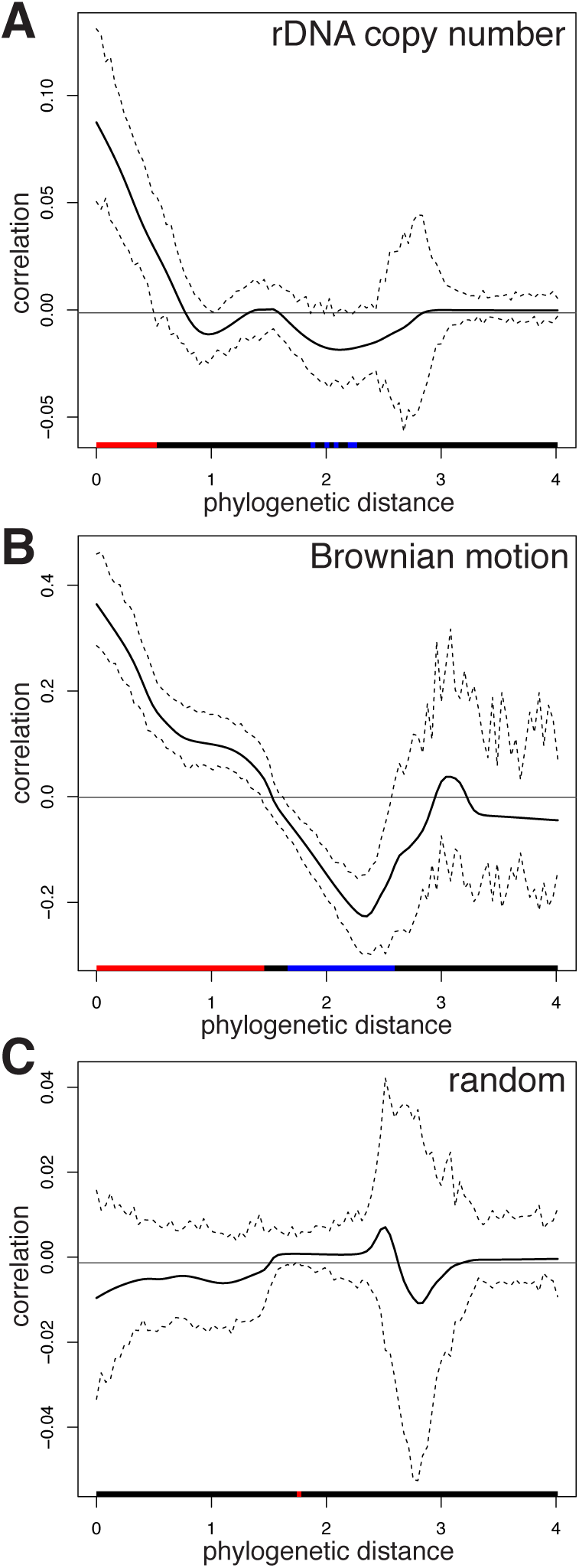
Phylocorrelograms of autocorrelation based on Moran’s *I*. Phylogenetic distance spatial autocorrelations between the SNP-based *S. cerevisiae* phylogeny and the rDNA copy number phylogeny (**A**), a Brownian motion phylogeny (**B**), and random data (**C**) are plotted. Red segments beneath each phylocorrelogram indicate significant positive autocorrelation; black no significant autocorrelation, and blue significant negative autocorrelation. Dotted lines indicate autocorrelation 95% confidence intervals. Significance is based on comparison to zero phylogenetic autocorrelation (horizontal black line at 0). Note the differences in the y-axis scale.

Another feature that might explain the distribution of rDNA copy numbers between *S. cerevisiae* populations is the environment, given that nutritional conditions have been proposed to influence copy number [65, 66]. To investigate this, we compared the rDNA copy numbers from two phylogenetically-diverged *S. cerevisiae* populations that are associated with oak trees, which we took as a proxy for similar environments. We found the oak populations did not show significantly different copy numbers (*p*-value = 0.52), as expected if environment is contributing to copy number. Thus, rDNA copy number might be partially determined by the environmental conditions the population has evolved in. However, we found no consistent pattern of similarities or differences with the copy numbers of the nearest phylogenetic neighbours of these oak clades (**Supplementary Information**), thus these results may simply represent stochastic variation. We suggest that a better understanding of what environmental factors modulate rDNA copy number is necessary before we can properly evaluate the impact of the environment on patterns of rDNA copy number variation.

Finally, we wondered whether large range in estimated *S. cerevisiae* rDNA copy number (22-227 copies) might reflect an unusually large variance in copy number in this species, given this range is almost the same as that reported across 91 different fungal species from three different fungal phyla (11-251 copies, excluding one outlier of 1442 copies; **Fig. 7**) [5]. However, comparing the *S. cerevisiae* copy number range generated by drawing twelve *S. cerevisiae* isolates at random from our data 1,000 times to that previously measured across twelve isolates of one fungal species (*Suillus brevipes*; [5]) shows that the *S. brevipes* range falls in the middle of the *S. cerevisiae* distribution of copy number ranges (**Fig 7**). These results suggest that *S. cerevisiae* rDNA copy number is no more variable than that of *S. brevipes* at least, and illustrate the tremendous inter-individual variation in rDNA copy number that is likely also the case for many other eukaryotic species.

**Figure 7.**
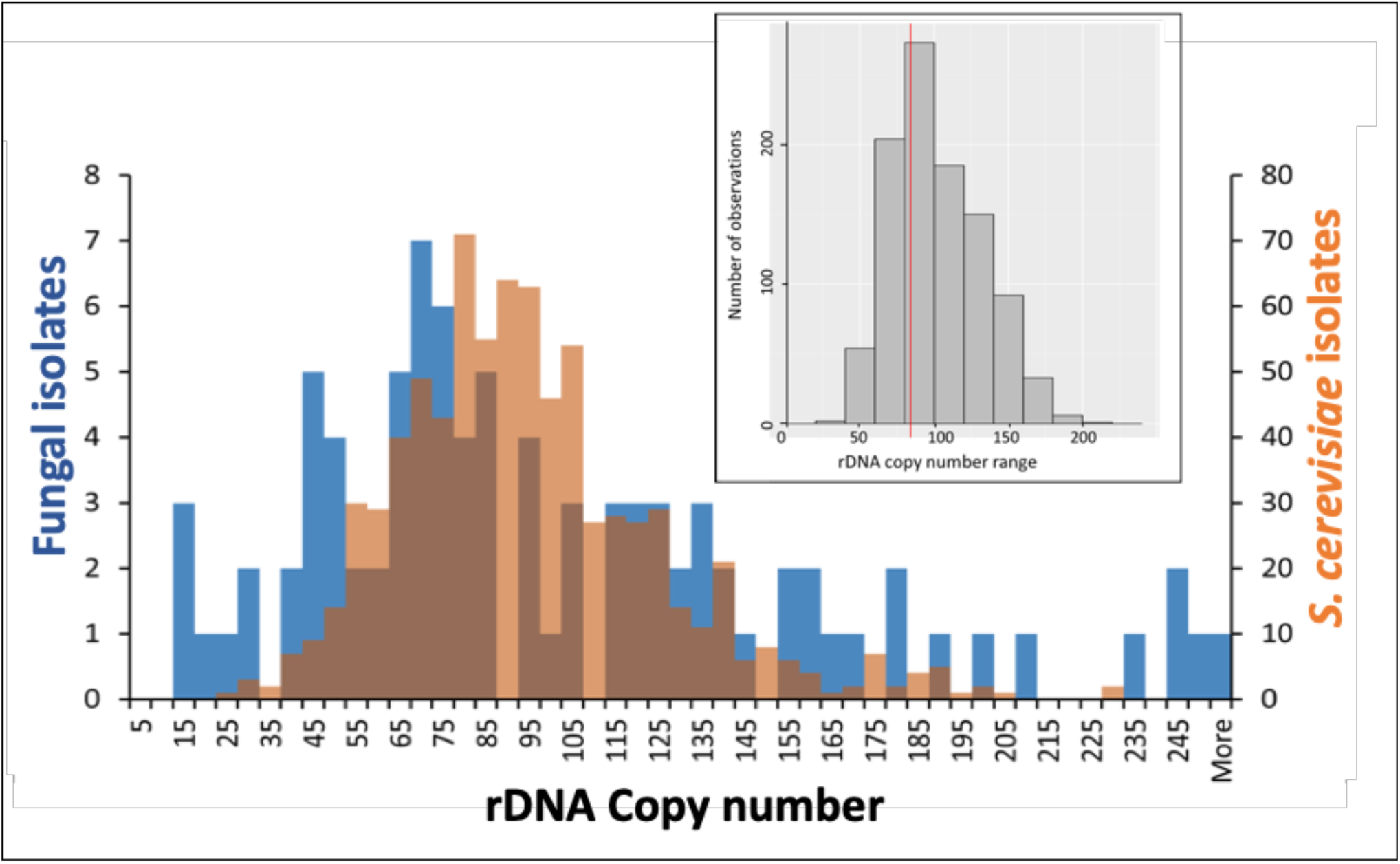
Distribution of rDNA copy number for fungal and *S. cerevisiae* isolates. The main histogram represents rDNA copy number (x-axis) for 91 previously published fungal taxa (blue bars, y-axis on left; [5]) and the 788 *S. cerevisiae* isolates (orange bars; y-axis on right) from this study. Brown represents overlaps. Inset histogram shows the distribution of total rDNA copy number ranges from 1,000 randomly drawn sets of twelve *S. cerevisiae* isolates. The red vertical line represents the total copy number range (84) observed amongst twelve *Suillus brevipes* isolates [5].

### Conclusions

Our results demonstrate that modal coverage can be used to robustly determine rDNA copy number from NGS data. Using our novel approach, we demonstrate that the mean rDNA copy number across all wild *S. cerevisiae* populations is 92. This is substantially lower than the copy numbers documented for lab *S. cerevisiae* strains, but overlaps the ‘typical’ rDNA copy numbers reported for fungi [5]. We show that *S. cerevisiae* populations have different homeostatic rDNA copy numbers, consistent with a previous study using a much smaller sample size [14]. We found some correlation between rDNA copy number and phylogeny, but not enough to suggest that homeostatic copy number is simply drifting apart with increasing phylogenetic distance. We also provide circumstantial evidence that environmental factors might help drive the homeostatic rDNA copy number differences. This is consistent with demonstrations that nutritional factors can induce physiological rDNA copy number changes [65, 66] and that such differences have phenotypic consequences [8, 12, 17, 22, 27–35]. However, it has been shown that rDNA copy number does not correlate with trophic mode in fungi [5] and we cannot exclude stochastic copy number variation explaining our environmental results. Therefore, more work is required to determine what really drives copy number dynamics between populations. One caveat to our conclusions is that while studies from a variety of organisms have demonstrated that copy number recovers from perturbation [7–10], presumably as a result of mechanisms maintaining homeostatic copy number [26], some recent studies in *S. cerevisiae* and *Drosophila* have reported the persistence of stochastic copy number changes without recovery [65, 67]. It will be important to reconcile these conflicting results and to determine to what extent the population-level differences we observe are the result of copy number homeostasis (as we interpret them) versus copy number inertia.

Our results showing population-level differences in rDNA copy number suggest that such differences can arise relatively quickly in evolutionary time, although the very high level of copy number variation between individuals acts to obscure this pattern. Therefore, it is important to take the large variances and rapid copy number dynamics of the rDNA into account when interpreting the impact of copy number variation on phenotype. Bioinformatics pipelines, such as the one we have developed here, in conjunction with the increasing availability of appropriate NGS datasets provide a way to establish baseline data on rDNA copy number variation between cells, individuals, populations, and species, as well as to investigate the phenotypic consequences of this variation. Finally, while we report population-level differences in rDNA copy number in *S. cerevisiae*, diverse human populations have been reported to not differ in rDNA copy number [12, 46]. Whether this reflects a difference in biology (such as differences in the level of genetic divergence between populations) or an incomplete understanding of human population rDNA copy number will require further clarification.

## Acknowledgements

We thank Gianni Liti (IRCAN, CNRS, INSERM, Université Côte d’Azur, University of Nice) for kindly providing strains. We acknowledge the University of Auckland Centre for eResearch (CER) and the New Zealand eScience Infrastructure (NeSI) for providing high performance computing facilities, consulting support, and training services. We thank Auckland Genomics for advice and whole genome sequencing, and Kevin Chang for help with statistical analyses. We thank Alastair Harris for helpful suggestions, and the Ganley Lab for helpful comments and feedback on the manuscript. This work was supported by a grant from the New Zealand Marsden Fund (14-MAU-053) and a University of Auckland Faculty Research Development Grant (3712288), both to ARDG.

## References

1. Long EO, Dawid IB. Repeated genes in eukaryotes. Annu Rev Biochem. 1980;49:727–64.

2. McStay B, Grummt I. The epigenetics of rRNA genes: From molecular to chromosome biology. Annu Rev Cell Dev Biol. 2008;24:131–57.

3. Prokopowich CD, Gregory TR, Crease TJ. The correlation between rDNA copy number and genome size in eukaryotes. Genome. 2003;46:48–50.

4. Torres-Machorro AL, Hernández R, Cevallos AM, López-Villasenor I. Ribosomal RNA genes in eukaryotic microorganisms: witnesses of phylogeny? FEMS Microbiol Rev. 2010;34:59–86.

5. Lofgren LA, Uehling JK, Branco S, Bruns TD, Martin F, Kennedy PG. Genome-based estimates of fungal rDNA copy number variation across phylogenetic scales and ecological lifestyles. Mol Ecol. 2019;28(4):721–30. doi: 10.1111/mec.14995.

6. Iida T, Kobayashi T. How do cells count multi-copy genes?: “Musical Chair” model for preserving the number of rDNA copies. Curr Genet. 2019;65(4):883–5. doi: 10.1007/s00294-019-00956-0.

7. Kobayashi T, Heck DJ, Nomura M, Horiuchi T. Expansion and contraction of ribosomal DNA repeats in *Saccharomyces cerevisiae*: requirement of replication fork blocking (Fob1) protein and the role of RNA polymerase I. Genes and Development. 1998;12:3821–30.

8. Hawley RS, Marcus CH. Recombinational controls of rDNA redundancy in *Drosophila*. Annu Rev Genet. 1989;23:87–120.

9. Russell PJ, Rodland KD. Magnification of rRNA gene number in a *Neurospora crassa* strain with a partial deletion of the nucleolus organizer. Chromosoma. 1986;93:337–40.

10. Rodland KD, Russell PJ. Regulation of ribosomal RNA cistron number in a strain of *Neurospora crassa* with a duplication of the nucleolus organizer region. Biochimica et Biophysica Acta. 1982;697:162–9.

11. Lyckegaard EM, Clark AG. Ribosomal DNA and Stellate gene copy number variation on the Y chromosome of *Drosophila melanogaster*. PNAS. 1989;86(6):1944–8. doi: 10.1073/pnas.86.6.1944.

12. Gibbons JG, Branco AT, Yu S, Lemos B. Ribosomal DNA copy number is coupled with gene expression variation and mitochondrial abundance in humans. Nature Communications. 2014;5:4850. doi: 10.1038/ncomms5850.

13. Cowen LE, Sanglard D, Calabrese D, Sirjusingh C, Anderson JB, Kohn LM. Evolution of drug resistance in experimental populations of *Candida albicans*. J Bacteriol. 2000;182:1515–22.

14. West C, James SA, Davey RP, Dicks J, Roberts IN. Ribosomal DNA sequence heterogeneity reflects intraspecies phylogenies and predicts genome structure in two contrasting yeast species. Syst Biol. 2014;63(4):543–54. doi: 10.1093/sysbio/syu019.

15. Herrera ML, Vallor AC, Gelfond JA, Patterson TF, Wickes BL. Strain-dependent variation in 18S ribosomal DNA Copy numbers in *Aspergillus fumigatus*. J Clin Microbiol. 2009;47(5):1325–32. doi: 10.1128/JCM.02073-08.

16. Stults DM, Killen MW, Pierce HH, Pierce AJ. Genomic architecture and inheritance of human ribosomal RNA gene clusters. Genome Res. 2008;18:13–8.

17. Ide S, Miyazaki T, Maki H, Kobayashi T. Abundance of ribosomal RNA gene copies maintains genome integrity. Science. 2010;327:693–6.

18. French SL, Osheim YN, Cioci F, Nomura M, Beyer AL. In exponentially growing *Saccharomyces cerevisiae* cells, rRNA synthesis is determined by the summed RNA polymerase I loading rate rather than the number of active genes. Mol Cell Biol. 2003;23:1558–68.

19. Kobayashi T, Ganley ARD. Recombination regulation by transcription-induced cohesin dissociation in rDNA repeats. Science. 2005;309:1581–4.

20. Szostak JW, Wu R. Unequal crossing over in the ribosomal DNA of *Saccharomyces cerevisiae*. Nature. 1980;284:426–30.

21. Ganley ARD, Kobayashi T. Monitoring the rate and dynamics of concerted evolution in the ribosomal DNA repeats of *Saccharomyces cerevisiae* using experimental evolution. Mol Biol Evol. 2011;28:2883–91.

22. Salim D, Gerton JL. Ribosomal DNA instability and genome adaptability. Chromosome Research. 2019;27(1-2):73–87. doi: 10.1007/s10577-018-9599-7.

23. Ganley ARD, Kobayashi T. Highly efficient concerted evolution in the ribosomal DNA repeats: total rDNA repeat variation revealed by whole-genome shotgun sequence data. Genome Res. 2007;17:184–91.

24. Eickbush TH, Eickbush DG. Finely orchestrated movements: evolution of the ribosomal RNA genes. Genetics. 2007;175:477–85.

25. James SA, O’Kelly MJT, Carter DM, Davey RP, van Oudenaarden A, Roberts IN. Repetitive sequence variation and dynamics in the ribosomal DNA array of *Saccharomyces cerevisiae* as revealed by whole-genome resequencing. Genome Res. 2009;19:626–35.

26. Iida T, Kobayashi T. RNA polymerase I activators count and adjust ribosomal RNA gene copy number. Mol Cell. 2019;73(4):645–54. doi: 10.1016/j.molcel.2018.11.029.

27. Delany ME, Muscarella DE, Bloom SE. Effects of rRNA gene copy number and nucleolar variation on early development: inhibition of gastrulation in rDNA-deficient chick embryos. J Hered. 1994;85(3):211–7. doi: 10.1093/oxfordjournals.jhered.a111437.

28. Kobayashi T. Regulation of ribosomal RNA gene copy number and its role in modulating genome integrity and evolutionary adaptibility in yeast. Cell Mol Life Sci. 2011;68:1395–403.

29. Paredes S, Maggert KA. Ribosomal DNA contributes to global chromatin regulation. PNAS. 2009;106:17829–34.

30. Paredes S, Branco AT, Hartl DL, Maggert KA, Lemos B. Ribosomal DNA deletions modulate genome-wide gene expression: “rDNA-sensitive” genes and natural variation. PLoS Genet. 2011;7:e1001376.

31. Michel AH, Kornmann B, Dubrana K, Shore D. Spontaneous rDNA copy number variation modulates Sir2 levels and epigenetic gene silencing. Genes and Development. 2005;19:1199–210.

32. Bughio F, Maggert KA. The peculiar genetics of the ribosomal DNA blurs the boundaries of transgenerational epigenetic inheritance. Chromosome Research. 2019;27(1-2):19–30. doi: 10.1007/s10577-018-9591-2.

33. Cullis CA. Quantitative variation of ribosomal RNA genes in flax genotrophs. Heredity. 1979;42:237–46.

34. Xu B, Li H, Perry JM, Singh VP, Unruh J, Yu Z, et al. Ribosomal DNA copy number loss and sequence variation in cancer. PLoS Genet. 2017;13(6):e1006771. doi: 10.1371/journal.pgen.1006771.

35. Zhou J, Sackton TB, Martinsen L, Lemos B, Eickbush TH, Hartl DL. Y chromosome mediates ribosomal DNA silencing and modulates the chromatin state in *Drosophila*. PNAS. 2012;109(25):9941–6. doi: 10.1073/pnas.1207367109.

36. Ritossa FM, Spiegelman S. Localization of DNA complementary to ribosomal RNA in the nucleolus organizer region of *Drosophila melanogaster*. PNAS. 1965;53:737–45. doi: 10.1073/pnas.53.4.737.

37. Wallace H, Birnstiel ML. Ribosomal cistrons and the nucleolar organizer. Biochimica et Biophysica Acta. 1966;114(2):296–310. doi: 10.1016/0005-2787(66)90311-x.

38. Schweizer E, MacKechnie C, Halvorson HO. The redundancy of ribosomal and transfer RNA genes in *Saccharomyces cerevisiae*. J Mol Biol. 1969;40:261–77.

39. Matsuda K, Siegel A. Hybridization of plant ribosomal RNA to DNA: the isolation of a DNA component rich in ribosomal RNA cistrons. PNAS. 1967;58(2):673–80. doi: 10.1073/pnas.58.2.673.

40. Maleszka R, Clark-Walker GD. Yeasts have a four-fold variation in ribosomal DNA copy number. Yeast. 1993;9:53–8.

41. Saka K, Takahashi A, Sasaki M, Kobayashi T. More than 10% of yeast genes are related to genome stability and influence cellular senescence via rDNA maintenance. Nucleic Acids Res. 2016;44(9):4211–21. doi: 10.1093/nar/gkw110.

42. Paredes S, Maggert KA. Expression of *I-Cre*I endonuclease generates deletions within the rDNA of Drosophila. Genetics. 2009;181:1661–71.

43. Chestkov IV, Jestkova EM, Ershova ES, Golimbet VE, Lezheiko TV, Kolesina NY, et al. Abundance of ribosomal RNA gene copies in the genomes of schizophrenia patients. Schizophrenia Research. 2018;197:305–14. doi: 10.1016/j.schres.2018.01.001.

44. LeRiche K, Eagle SH, Crease TJ. Copy number of the transposon, *Pokey*, in rDNA is positively correlated with rDNA copy number in *Daphnia obtuse*. PLoS One. 2014;9(12):e114773. doi: 10.1371/journal.pone.0114773.

45. Son J, Hannan KM, Poortinga G, Hein N, Cameron DP, Ganley ARD, et al. rDNA chromatin activity status as a biomarker of sensitivity to the RNA polymerase I transcription inhibitor CX-5461. Frontiers in Cell and Developmental Biology. 2020;8:568.

46. Valori V, Tus K, Laukaitis C, Harris DT, LeBeau L, Maggert KA. Human rDNA copy number is unstable in metastatic breast cancers. Epigenetics. 2020;15(1-2):85–106. doi: 10.1080/15592294.2019.1649930.

47. Alanio A, Sturny-Leclere A, Benabou M, Guigue N, Bretagne S. Variation in copy number of the 28S rDNA of *Aspergillus fumigatus* measured by droplet digital PCR and analog quantitative real-time PCR. J Microbiol Methods. 2016;127:160–3. doi: 10.1016/j.mimet.2016.06.015.

48. Salim D, Bradford WD, Freeland A, Cady G, Wang J, Pruitt SC, et al. DNA replication stress restricts ribosomal DNA copy number. PLoS Genet. 2017;13(9):e1007006. doi: 10.1371/journal.pgen.1007006.

49. Rosato M, Kovarik A, Garilleti R, Rossello JA. Conserved organisation of 45S rDNA sites and rDNA gene copy number among major clades of early land plants. PLoS One. 2016;11(9):e0162544. doi: 10.1371/journal.pone.0162544.

50. Xu J, Xu Y, Yonezawa T, Li L, Hasegawa M, Lu F, et al. Polymorphism and evolution of ribosomal DNA in tea (*Camellia sinensis*, Theaceae). Mol Phylogen Evol. 2015;89:63–72. doi: 10.1016/j.ympev.2015.03.020.

51. Xu J, Zhang Q, Xu X, Wang Z, Qi J. Intragenomic variability and pseudogenes of ribosomal DNA in stone flounder *Kareius bicoloratus*. Mol Phylogen Evol. 2009;52(1):157–66. doi: 10.1016/j.ympev.2009.03.031.

52. Agrawal S, Ganley ARD. Complete sequence construction of the highly repetitive ribosomal RNA gene repeats in eukaryotes using whole genome sequence data. Methods in Molecular Biology. 2016;1455:161–81. doi: 10.1007/978-1-4939-3792-9_13

53. Buckler ES, Ippolito A, Holtsford TP. The evolution of ribosomal DNA: Divergent paralogues and phylogenetic implications. Genetics. 1997;145:821–32.

54. Mayol M, Rosselló JA. Why nuclear ribosomal DNA spacers (ITS) tell different stories in *Quercus*. Mol Phylogen Evol. 2001;19:167–76.

55. Wang M, Lemos B. Ribosomal DNA copy number amplification and loss in human cancers is linked to tumor genetic context, nucleolus activity, and proliferation. PLoS Genet. 2017;13(9):e1006994. doi: 10.1371/journal.pgen.1006994.

56. Gong W, Marchetti A. Estimation of 18S gene copy number in marine eukaryotic plankton using a next-generation sequencing approach. Frontiers in Marine Science. 2019;6:219.

57. Cubillos FA, Louis EJ, Liti G. Generation of a large set of genetically tractable haploid and diploid *Saccharomyces* strains. FEMS Yeast Res. 2009;9(8):1217–25. doi: 10.1111/j.1567-1364.2009.00583.x.

58. Liti G, Carter DM, Moses AM, Warringer J, Parts L, James SA, et al. Population genomics of domestic and wild yeasts. Nature. 2009;458(7236):337–41. doi: 10.1038/nature07743.

59. Cox MP, Peterson DA, Biggs PJ. SolexaQA: At-a-glance quality assessment of Illumina second-generation sequencing data. BMC Bioinformatics. 2010;11:485. doi: 10.1186/1471-2105-11-485.

60. Kumar S, Stecher G, Li M, Knyaz C, Tamura K. MEGA X: Molecular evolutionary genetics analysis across computing platforms. Mol Biol Evol. 2018;35(6):1547–9. doi: 10.1093/molbev/msy096.

61. Keck F, Rimet F, Bouchez A, Franc A. phylosignal: an R package to measure, test, and explore the phylogenetic signal. Ecology and Evolution. 2016;6(9):2774–80. doi: 10.1002/ece3.2051.

62. Pennell MW, Eastman JM, Slater GJ, Brown JW, Uyeda JC, FitzJohn RG, et al. geiger v2.0: an expanded suite of methods for fitting macroevolutionary models to phylogenetic trees. Bioinformatics. 2014;30(15):2216–8. doi: 10.1093/bioinformatics/btu181.

63. Yoon S, Xuan Z, Makarov V, Ye K, Sebat J. Sensitive and accurate detection of copy number variants using read depth of coverage. Genome Res. 2009;19(9):1586–92. doi: 10.1101/gr.092981.109.

64. Peter J, De Chiara M, Friedrich A, Yue JX, Pflieger D, Bergstrom A, et al. Genome evolution across 1,011 *Saccharomyces cerevisiae* isolates. Nature. 2018;556(7701):339–44. Epub 2018/04/13. doi: 10.1038/s41586-018-0030-5.

65. Aldrich JC, Maggert KA. Transgenerational inheritance of diet-induced genome rearrangements in Drosophila. PLoS Genet. 2015;11(4):e1005148. Epub 2015/04/18. doi: 10.1371/journal.pgen.1005148.

66. Jack CV, Cruz C, Hull RM, Keller MA, Ralser M, Houseley J. Regulation of ribosomal DNA amplification by the TOR pathway. PNAS. 2015;112(31):9674–9. Epub 2015/07/22. doi: 10.1073/pnas.1505015112.

67. Mansisidor A, Molinar T, Srivastava P, Dartis DD, Pino Delgado A, Blitzblau HG, et al. Genomic copy-number loss is rescued by self-limiting production of DNA circles. Mol Cell. 2018;72(3):583–93. Epub 2018/10/09. doi: 10.1016/j.molcel.2018.08.036.

