## Supplementary Figure 1 for "A new method for determining ribosomal DNA copy number shows differences between *Saccharomyces cerevisiae* populations"

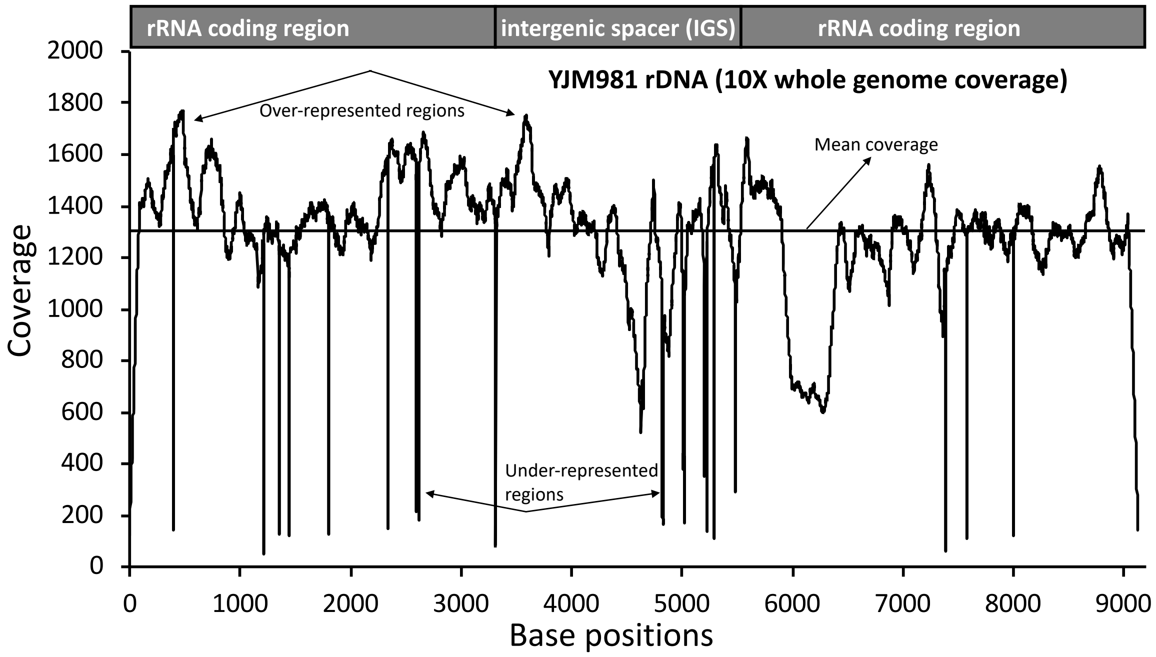


**Supplementary Figure 1. Sequence read coverage across the rDNA for a *S. cerevisiae* isolate.** Coverage (Y-axis) is plotted against rDNA base position (X-axis). The black horizontal line depicts the mean coverage value. The rRNA coding and intergenic spacer regions are indicated above. Regions over- and under-represented for coverage are indicated. The coverage data were generated by aligning reads from *S. cerevisiae* isolate YJM981 (from the 1002 Yeast Genome project; Peter et al. 2018) to the yeast genome with rDNA reference used in this study.
