## Supplementary Figure 2 for "A new method for determining ribosomal DNA copy number shows differences between *Saccharomyces cerevisiae* populations"

| | R1-1 | R1-2 | R2-1 | R2-2 | R3-1 | R3-2 |
|---|---|---|---|---|---|---|
| **WG1-1** | CN1 | CN2 | CN3 | CN4 | CN5 | CN6 |
| **WG1-2** | CN7 | CN8 | CN9 | CN10 | CN11 | CN12 |
| **WG2-1** | CN13 | CN14 | CN15 | CN16 | CN17 | CN18 |
| **WG2-2** | CN19 | CN20 | CN21 | CN22 | CN23 | CN24 |
| **WG3-1** | CN25 | CN26 | CN27 | CN28 | CN29 | CN30 |
| **WG3-2** | CN31 | CN32 | CN33 | CN34 | CN35 | CN36 |

**Supplementary Figure 2. Schematic showing how rDNA copy number is calculated from the three highest frequency coverage bins from both the whole genome and the rDNA.** The three highest (peak) coverage bins for both rDNA and whole genome are determined. R represents the three highest peak values for rDNA, WG the three highest peak values for whole genome. The -1 and -2 designations refer to the upper and lower values of the coverage bin range, respectively. This produces a total of 36 ratios of values, as indicated by CN. The estimated copy number is then the mean of these 36 CN values.
