## Supplementary Figure 3 for "A new method for determining ribosomal DNA copy number shows differences between *Saccharomyces cerevisiae* populations"

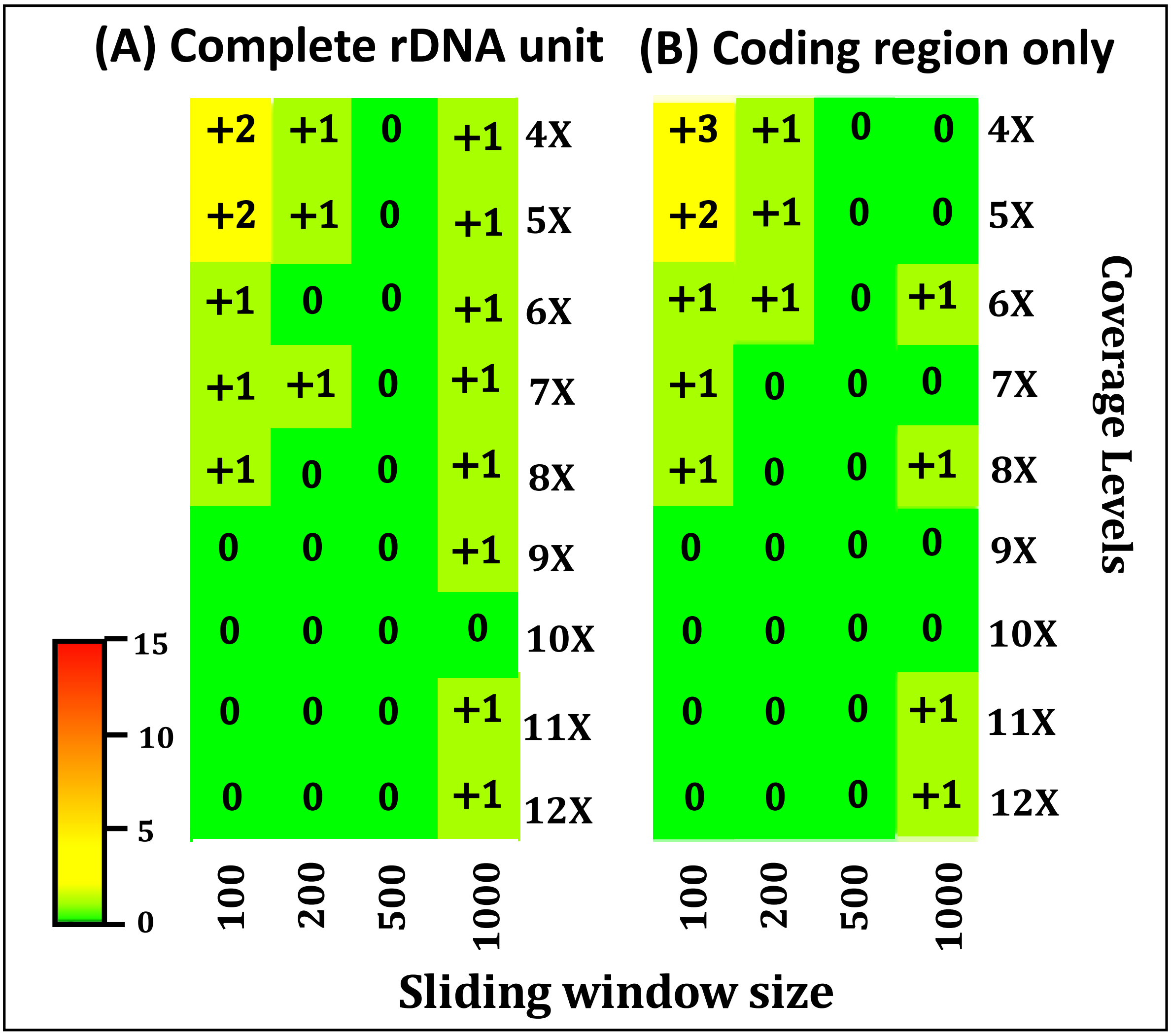


**Supplementary Figure 3. Copy number estimates are similar when only using the rRNA coding region.** Each cell represents the deviation of the calculated modal rDNA copy number from 20 for each coverage level and sliding window size combination. (**A**) uses coverage information from the whole rDNA repeat, while (**B**) only uses the information from the rRNA coding region. The heatmap scale used is indicated and deviations are rounded to the nearest integer.
