## Supplementary Figure 4 for "A new method for determining ribosomal DNA copy number shows differences between *Saccharomyces cerevisiae* populations"

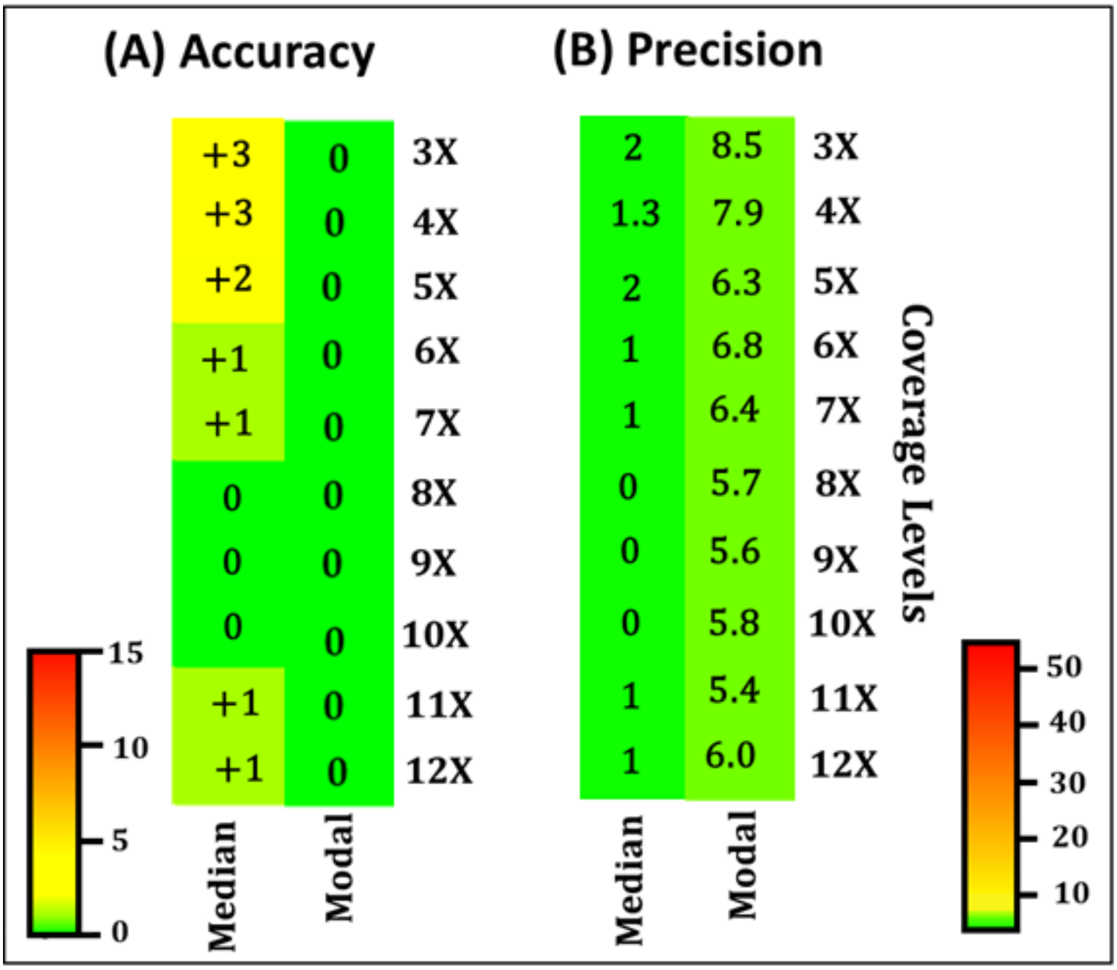


**Supplementary Figure 4. Accuracy and precision of the median coverage approach versus the modal coverage approach.** Each cell represents the deviation of the estimated mean rDNA copy number, calculated from the 100 technical replicates for each coverage level, from the "true" rDNA copy number (20), using a sliding window size of 600 bp. The heatmap scales used are indicated, and rDNA copy number was rounded to the nearest integer.
