## Supplementary Figure 5 for "A new method for determining ribosomal DNA copy number shows differences between *Saccharomyces cerevisiae* populations"

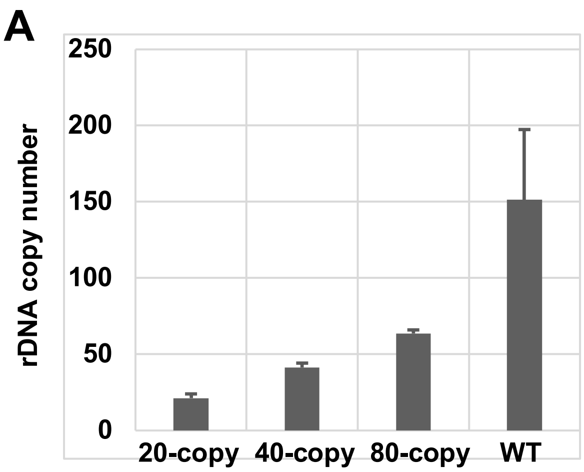


| Strain | 20-copy | 40-copy | 80-copy | WT |
|---|---|---|---|---|
| rDNA copy number | 21.23 | 41.36 | 63.61 | 151.33 |
| SD | 2.77 | 2.84 | 2.37 | 46.09 |

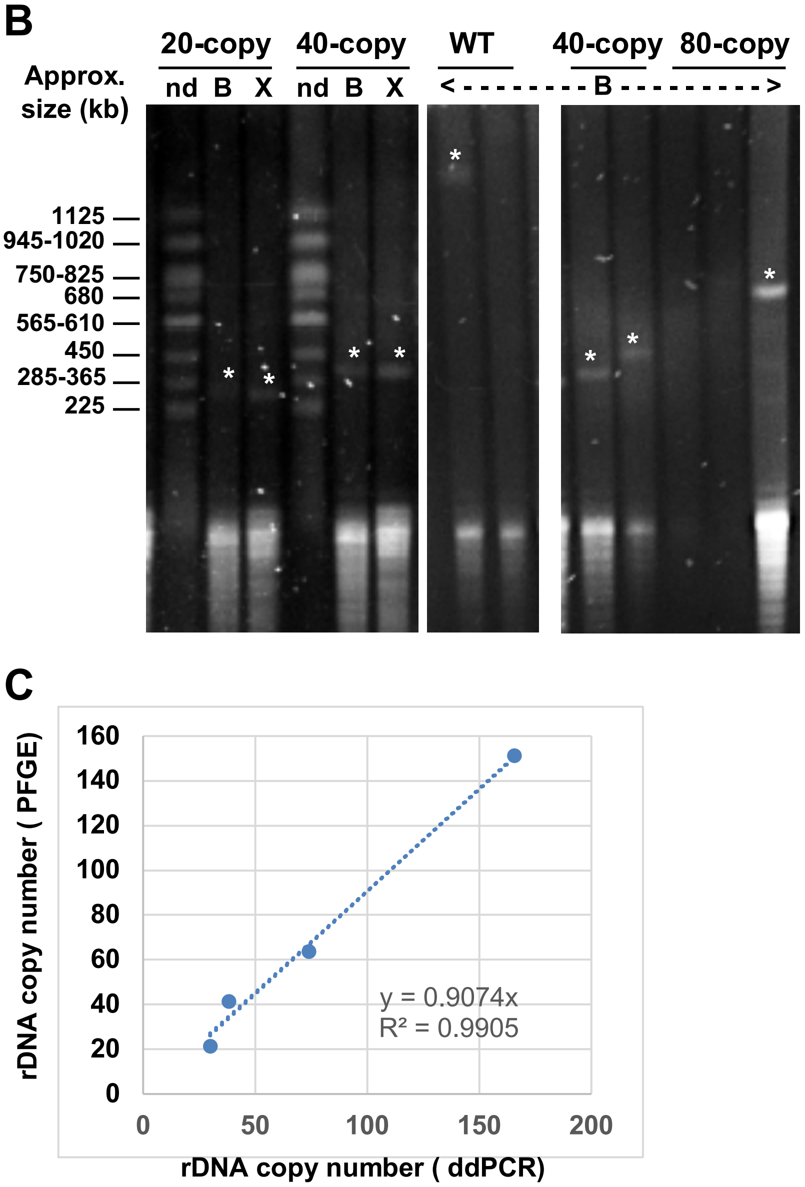

**Supplementary Figure 5. rDNA copy number measurement by droplet digital PCR (ddPCR) and pulsed field gel electrophoresis (PFGE).** (**A**) ddPCR estimates of rDNA copy number for the indicated laboratory strains using at least three independent biological replicates. Mean values and standard deviations (SD) are given in the table below. (**B**) Ethidium bromide-stained PFG showing two independent experiments detecting the undigested chromosome profile (nd) and the rDNA array (white stars) for the indicated strains. Entire rDNA arrays are detected after enzymatic restriction with *Xho*I (X) or *BamHI* (B) enzymes that do not cut within the rDNA array. Sizes of undigested *S. cerevisiae* chromosomes were used to evaluate each rDNA array size. (**C**) Correlation between rDNA copy number evaluated by PFGE and ddPCR for the four strains.
