## Supplementary Figure 6 for "A new method for determining ribosomal DNA copy number shows differences between *Saccharomyces cerevisiae* populations"

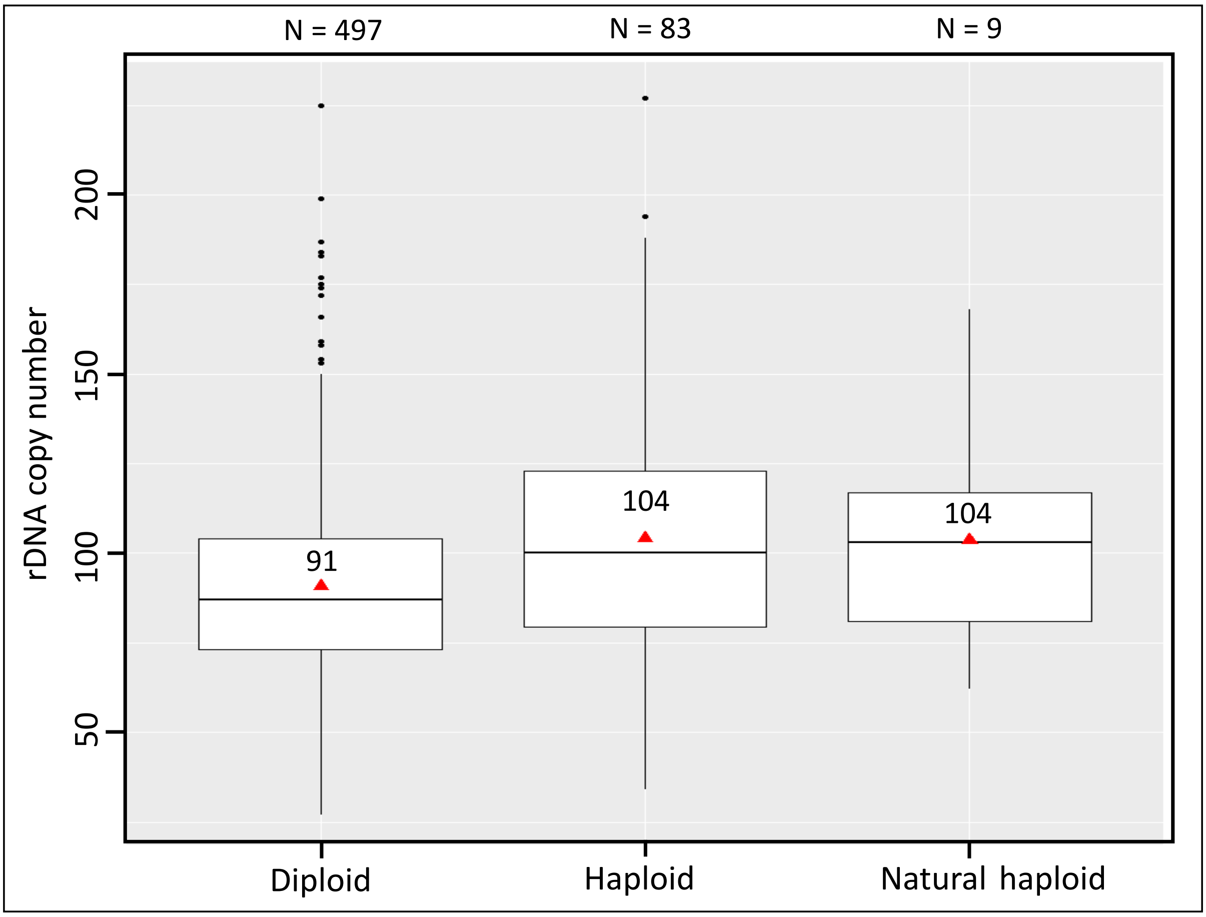


**Supplementary Figure 7. Ploidy difference does not explain the higher rDNA copy numbers in lab versus wild isolates.** Mean copy numbers of all 1002 Yeast Genome project strains indicated as purely diploid or purely haploid from Peter et al (2018) are plotted as a box plot. The haploid isolates are further divided into those naturally found as haploids (natural haploid) versus those subsequently converted to haploid post-isolation. The number of isolates are indicated at the top, medians are indicated by the horizontal lines in the boxes, and means by red triangles with the values shown above. While haploid rDNA copy numbers are slightly higher than diploid copy numbers, the magnitude is not sufficient to explain the difference between lab strains and wild isolates.
