## Supplementary information for "A new method for determining ribosomal DNA copy number shows differences between *Saccharomyces cerevisiae* populations"

**rDNA copy number differences are not explained by ploidy differences**

The ANOVA results showing population-level differences in rDNA copy number are consistent with previous results showing laboratory *S. cerevisiae* strains typically have higher (150-250) homeostatic copy numbers (e.g. Kobayashi et al. 1998; West et al. 2014; Salim et al. 2017) than those we estimated for most *S. cerevisiae* wild isolates in this study. However, an alternative explanation is that copy number is influenced by ploidy, as the lab strains are haploid, while most isolates in the 1002 Yeast Genome project are diploid (Peter et al. 2018). For example, if rDNA copy number is determined on a per cell rather than per genome basis, we would expect haploid strains to have a higher (approximately double) per genome copy number than diploid strains. To test this, we calculated the mean rDNA copy number for all 83 isolates identified as haploid, and all 497 strains identified as diploid, from the 1002 Yeast Genome data. While we found a trend in this direction (**Supplementary Figure 6**; haploid copy number mean of ~104 versus ~91 for diploids), this is not sufficient to explain the copy number differences between lab strains and the 1002 Yeast Genome wild isolates. This conclusion holds even when we restrict our analysis of haploids to just the nine naturally-occurring haploids defined by Peter et al. (2018), as these have the same mean copy number as all haploids (**Supplementary Figure 6**).

**Correlation of rDNA copy number with environment**

Our results suggest that different *S. cerevisiae* populations have different homeostatic rDNA copy numbers, but phylogeny does not appear to fully explain the distribution of rDNA copy numbers. It is possible that copy number evolves in response to different environmental conditions in a way that is not completed correlated with phylogeny. To look for evidence of this, we compared the rDNA copy numbers of phylogenetically divergent populations that share similar environments, in this case association with oak (a Mediterranean oak population and a North American oak population). We also compared the copy numbers of these populations to those of their nearest phylogenetic neighbours. Two independent statistical tests showed no significant difference in rDNA copy number between populations from similar environments (Welch two sample t-test *p*-value = 0.52; Mann Whitney Wilcoxon rank sum test *p*-value = 0.38), as expected if environment rather than the ancestry is driving rDNA copy numbers. Therefore, the results of this analysis are consistent with environment helping to drive the rDNA copy number patterns we observe. However, these analyses are severely limited by our lack of knowledge of what, if any, environmental factor(s) drive differences in rDNA copy number.

**References**

Kobayashi T, Heck DJ, Nomura M, Horiuchi T. 1998. Expansion and contraction of ribosomal DNA repeats in *Saccharomyces cerevisiae*: requirement of replication fork blocking (Fob1) protein and the role of RNA polymerase I. *Genes and Development* **12**: 3821-3830.

Lofgren LA, Uehling JK, Branco S, Bruns TD, Martin F, Kennedy PG. 2019. Genome-based estimates of fungal rDNA copy number variation across phylogenetic scales and ecological lifestyles. *Mol Ecol* **28**: 721-730.

Peter J, De Chiara M, Friedrich A, Yue JX, Pflieger D, Bergstrom A, Sigwalt A, Barre B, Freel K, Llored A et al. 2018. Genome evolution across 1,011 *Saccharomyces cerevisiae* isolates. *Nature* **556**: 339-344.

Salim D, Bradford WD, Freeland A, Cady G, Wang J, Pruitt SC, Gerton JL. 2017. DNA replication stress restricts ribosomal DNA copy number. *PLoS Genet* **13**: e1007006.

West C, James SA, Davey RP, Dicks J, Roberts IN. 2014. Ribosomal DNA sequence heterogeneity reflects intraspecies phylogenies and predicts genome structure in two contrasting yeast species. *Syst Biol* **63**: 543-554.
