## Supplementary Table 1 for "A new method for determining ribosomal DNA copy number shows differences between *Saccharomyces cerevisiae* populations"

**Supplementary Table 1. Details of whole genome sequencing of the *S. cerevisiae* rDNA copy number strains**

| Strain | Number of reads | Sequence length | Library | Sequencer |
| --- | --- | --- | --- | --- |
| 20-copy | 1,232,762 | 150 bp paired end | Thruplex DNA-seq | MiSeq |
| 40-copy | 1,043,748 | 150 bp paired end | Thruplex DNA-seq | MiSeq |
| 80-copy | 1,060,342 | 150 bp paired end | Thruplex DNA-seq | MiSeq |
| WT-copy | 1,706,686 | 250 bp paired end | TruSeq LT | MiSeq |
| US-05 |  |  |  |  |
