## Supplementary Table 2 for "A new method for determining ribosomal DNA copy number shows differences between *Saccharomyces cerevisiae* populations"

**Supplementary Table 2. Comparisons of rDNA copy number estimation between methods**

| Sample | ddPCR rDNA copy number | Modal rDNA copy number | Mean rDNA copy number |
| --- | --- | --- | --- |
| WT | 151 | 157 | 133 |
| 20-copy | 21 | 20 | 18 |
| 40-copy | 41 | 36 | 32 |
| 80-copy | 64 | 64 | 57 |
