## Supplementary Table 4 for "A new method for determining ribosomal DNA copy number shows differences between *Saccharomyces cerevisiae* populations"

**Supplementary Table 4. Comparisons of rDNA copy numbers for 11 isolates estimated in this and previous studies**

| Strains | Previous mean coverage rDNA CN estimates | | rDNA CN estimates using modal coverage approach^a^ | rDNA CN estimates using mean coverage approach^b^ |
| --- | --- | --- | --- | --- |
| DBVPG 1788 | 75^c^ | 67^d^ | 87 | 85 |
| DBVPG 1106 | 112 | 98 | 110 | 110 |
| DBVPG 6044 | 120 | 107 | 131 | 139 |
| YJM981^e^ | 511 | 354 | 171 | 159 |
| UWO0S03-461-4 | 98 | 89 | 106 | 107 |
| W303 | 217 | 182 | 157 | 179 |
| Y12 | 85 | 78 | 105 | 103 |
| Y55 | 86 | 72 | 107 | 106 |
| Y9 | 88 | 79 | 188 | 178 |
| K11 | 54 | 50 | 138 | 106 |
| YPS128 | 64 | 62 | 89 | 89 |

^a^ Copy number estimates using modal coverage method were performed using 10-fold whole genome sequence coverage and a sliding window of 600 bp.

^b^ Copy number estimates using the mean coverage approach were performed using 10-fold whole genome sequence coverage.

^c^ rDNA copy number estimates taken from James et al. (2009).

^d^ rDNA copy number estimates taken from West et al. (2014).

^e^ Previous estimates for this isolate were anomalously high. We are unsure why this is, given that our estimate, using both mean and modal coverage approaches, are within the “normal” range and the rDNA coverage does not look anomalous (**Supplementary Figure 1**).
