## Supplementary Table 5 for "A new method for determining ribosomal DNA copy number shows differences between *Saccharomyces cerevisiae* populations"

**Supplementary Table 5: Experimental testing of homeostatic rDNA copy number for wild and laboratory *S. cerevisiae* strains**

| **Strains/ isolates** | **Ploidy** | **Origin/ Background** | **rDNA CN**  **(James et al. 2009)** | **rDNA CN**  **(West et al. 2014)** | **rDNA CN**  **mean coverage** | **rDNA CN**  **modal coverage** | **rDNA CN**  **ddPCR experiment** | | |
| --- | --- | --- | --- | --- | --- | --- | --- | --- | --- |
|  |  |  |  |  |  |  | 15^b^ | 60^b^ mean^c^ | 60 |
| WT^a^ | n | Laboratory strain W303 | 217 | 182 | 179 | 157 | 213 | 185 | 130 |
|  |  |  |  |  |  |  |  |  | 217 |
|  |  |  |  |  |  |  |  |  | 208 |
| YJM981  Mat**a** derivative^e^ | n | Human clinical,  Italy  Cubillos et al. 2009 | 511 | 354 | 159 | 171 | 174 | 175 | 120 |
|  |  |  |  |  |  |  |  |  | 183 |
|  |  |  |  |  |  |  |  |  | 221 |
| DBVPG1373  Mat**a** derivative | n | Soil, Netherlands  Cubillos et al. 2009 | 80 | 75 | 57 | 78 | 69 | 85 | 77 |
|  |  |  |  |  |  |  |  |  | 72 |
|  |  |  |  |  |  |  |  |  | 107 |
| UWOPS03-461-4  (1005) | 2n | Nectar, Malaysia  Liti et al. 2009 | 98 | 89 | 107 | 106 | 85 | 95 | 113 |
|  |  |  |  |  |  |  |  |  | 88 |
|  |  |  |  |  |  |  |  |  | 83 |
| UWOPS03-461-4 Mat**a** derivative | n | Nectar, Malaysia  Cubillos et al. 2009 | ND | ND | ND | ND | 244 | 146 | 164 |
|  |  |  |  |  |  |  |  |  | 167 |
|  |  |  |  |  |  |  |  |  | 106 |
| UWOPS03-461-4  Matα derivative | n | Nectar, Malaysia  Cubillos et al. 2009 | ND | ND | ND | ND | ND | 109 | 108 |
|  |  |  |  |  |  |  |  |  | 115 |
|  |  |  |  |  |  |  |  |  | 105 |
| YPS128 | 2n | *Quercus alba* soil, USA  Liti et al. 2009 | 64 | 62 | 89 | 89 | 89 | 79 | 87 |
|  |  |  |  |  |  |  |  |  | 73 |
|  |  |  |  |  |  |  |  |  | 77 |
| DBVPG1788 | n | Vineyard soil, Finland  Liti et al. 2009 | 75 | 67 | 85 | 87 | 95 | 108 | 126 |
|  |  |  |  |  |  |  |  |  | 100 |
|  |  |  |  |  |  |  |  |  | 97 |

^a^ Laboratory strain

^b^ Number of generations before cell harvesting

^c^ The mean of the three values obtained for the three biological replicates in the right hand column

^d^ ND: Not Determined

^e^ Wild isolates were chosen to represent the range of rDNA copy numbers estimated during this work
